# Investigation of the effect of physiological factors on resting-state and task-based functional connectivity

**DOI:** 10.1101/2024.09.29.615715

**Authors:** S. Emad Askarinejad, Jean-Baptiste Poline, Georgios D. Mitsis

## Abstract

Understanding the brain’s functional network through functional connectivity (FC) is crucial for gaining deeper insights into brain functional mechanism and identifying a potential biomarker for diagnosing neurological disorders. Despite the development of various FC measures, their reliability under different conditions remains under-explored. Moreover, physiological noise can obscure true neural activity, and accordingly, introduce errors into FC patterns. This issue necessitates further investigation. In this study, we evaluate and compare the performance of various methods using Local Field Potential and Blood-Oxygen-Level-Dependent signals across different conditions. We also examine the impact of physiological artifacts on BOLD-FC results. Our comprehensive assessment covers multiple modalities of brain signals, diverse task paradigms, and varying noise levels. Our findings reveal that while Granger Causality-based methods exhibit significant limitations, particularly with BOLD data, multivariate techniques (e.g. partial correlation) demonstrate greater robustness in distinguishing between different types of connections within the network. Notably, our results indicate that physiological artifacts substantially affect FC values, leading to erroneous connectivity estimates, especially with bivariate methods. This research offers a foundational analysis of the effects of physiological artifacts on FC results and provides valuable insights for future studies.

## 1 Introduction

The study of brain function begins with two foundational concepts: functional specialization and functional integration. Functional specialization emphasizes the role of distinct brain regions in processing specific types of information. Numerous neuroimaging studies have adopted this perspective, identifying cortical regions that are predominantly modulated by particular stimuli. However, it is understood that executing a single function typically requires the involvement of multiple regions operating as nodes within a distributed network, a concept referred to as functional integration [1]. Functional integration encompasses two closely related branches−functional connectivity (FC) and effective connectivity (EC)−which, to some extent, share the same information about the brain’s functional network. FC is mathematically defined as the temporal dependency among spatially distinct brain regions [2], while EC is defined as the causal influence that one region (as a single node in a neural network) exerts over another [3]. Consequently, FC has emerged as a crucial measure for gaining a deeper understanding of brain function and as a biomarker with significant potential [4, 5].

Functional connectivity (FC) can be studied under two primary conditions: resting state and task execution. FC is most commonly investigated during wakeful rest (rest-FC), where the subject is not engaged in any specific task or exposed to external stimuli. Conversely, task-based FC (task-FC) examines the brain’s functional networks while tasks are being performed or external stimuli are present. Different tasks can alter FC patterns of the brain over multiple temporal scales, ranging from milliseconds to changes occurring over minutes, at both neural [6] and system [7] level. Several key similarities and differences between task-FC and rest-FC have been identified. For instance, during task performance, across-network connectivity often changes, with task-related regions showing increased functional connectivity and higher overall integration [8]. Additionally, the frequency profile of inter-regional communication is altered by task engagement, leading to a more balanced contribution of different frequencies to task-FC [9].

A variety of methods have been developed to assess functional connectivity (FC), aiming to determine both direct and indirect connections, as well as to reveal linear or nonlinear relationships between brain regions. These methods can analyze connections between two (*bivariate*) or multiple (*multivariate*) brain regions using amplitude or phase values in either the time or frequency domain. FC assessment techniques can be broadly classified into four main categories: 1) correlation or coherence-based methods, 2) phase synchronization measures, 3) information-based measures, and 4) Granger causality measures. As these methods are applied to the entire length of the recorded signals, which are assumed to be stationary, they are commonly referred to as static-FC techniques. In 2014, van Mierlo et al. [10] reviewed the performance of various FC assessment methods in predicting seizures and localizing the epileptogenic zone using scalp EEG signals, demonstrating their potential in this area. Subsequently, Mitsis et al. (2020) [11] applied FC measures to windowed, long-duration scalp EEG signals for seizure prediction, suggesting that longer recordings could enhance the robustness of seizure detection. Moreover, several studies have investigated the capability of different FC measures to map network structures. For instance, Winterhalder et al. (2005) [12] evaluated the effectiveness of various multivariate linear techniques in identifying direct interactions within linear, nonlinear, and non-stationary multivariate neural systems. Additionally, Smith et al. (2011) [13] conducted a comprehensive comparison of multiple FC measures in modeling network structures using simulated fMRI data, assessing their accuracy in identifying true network connections.

In this context, electrophysiological methods like electroencephalography (EEG) and magnetoencephalography (MEG), along with functional imaging techniques such as positron emission tomography (PET) and functional magnetic resonance imaging (fMRI), are widely used to characterize functional relationships in the brain. Each modality offers a complementary perspective on brain activity based on its unique properties. For instance, EEG captures the cortical electrical activity, providing high temporal resolution but low spatial resolution. In contrast, fMRI, measuring blood-oxygen-level-dependent (BOLD) signals, offers lower temporal resolution (on the order of seconds) but higher spatial resolution. Additionally, while EEG is recorded at the scalp level, fMRI provides insights into deeper brain regions by detecting changes in oxyhemoglobin and deoxyhemoglobin concentrations associated with neural activity.

Regarding noise contamination, since EEG signals are recorded at the scalp level, it must traverse various tissues which introduces multiple sources of noise. On the other hand, while fMRI is not subjected to the same types of noise, it is susceptible to physiological processes such as respiration and cardiac activity, which can contaminate BOLD-fMRI signals by inducing physiological artifacts. These artifacts can, in turn, affect the corresponding FC results, though the full extent of their impact is still not fully understood and is an area of ongoing discussion and investigation. In our previous study [14], we explored this issue by employing two commonly used FC methods: full correlation and partial correlation. We investigated the influence of physiological artifacts on the computed FC patterns during both rest and task conditions separately. The results indicated that partial correlation is less susceptible to these artifacts, leading to lower error values compared to full correlation. Moreover, in general, FC values computed during task execution exhibited lower error rates compared to those computed during the rest period.

In this study, building on our previous work, we employed several FC assessment measures, each offering a distinct perspective on brain network modeling, to evaluate their effectiveness in accurately identifying direct, indirect, and functional connections. Additionally, we assessed the robustness of these measures against the induced physiological artifacts and quantitatively analyzed the extent of their impact. To begin with, we conducted a realistic simulation to generate BOLD and local field potential (LFP) signals during both task and rest conditions, as detailed in Section 2. Furthermore, we explored the effects of physiological tasks, where artifacts are modulated by stimulus, as well as non-physiological tasks. Section 3 provides definitions and explanations of the FC measures utilized in this study. The main findings, along with their implications, are presented and discussed in Section 4. Finally, Section 5 concludes the paper by interpreting the findings, emphasizing their significance, and suggesting directions for future research.

## 2 Methods: Simulation

The brain functions as a dynamic network, with its various regions acting as interconnected nodes of the network. In this study, we utilized “*The V irtual Brain*” (TVB) [15], to generate different modalities of brain signals. TVB is an open-source simulation tool that can generate signals that closely mimic human brain activity. Initially, we simulated local field potentials (LFPs), which are electrical signals generated in the extracellular space surrounding neurons. Unlike electroencephalogram (EEG) signals, which are recorded at the scalp’s surface, LFPs are captured from deeper cortical tissues; however, both LFP and EEG exhibit similar oscillatory patterns. Subsequently, the simulated LFPs were fed into the balloon model and convolved with a hemodynamic response function (HRF) to produce blood-oxygen-level-dependent (BOLD) signals. To better reflect real-world scenarios in the simulation, we modeled physiological processes such as respiration and cardiac signals under various scenarios. Finally, to generate the physiological artifacts typically present in BOLD signals, the physiological signals were convolved with their corresponding physiological response functions (PRFs) and incorporated into the simulated BOLD signals. The following sections will provide detailed explanations of the simulation process for LFPs, BOLD, and physiological signals.

### 2.1 Local field potential (LFP)

TVB utilizes neural mass models to capture the behavior of large neuron populations. These models provide a mathematical framework that represents the collective activity of interconnected neurons within a network, rather than focusing on individual units. In TVB, networks can be structured at two distinct spatial scales: surface-based and region-based. For this study, we adapted the region-based spatial scale for our simulations. In this configuration, each brain region is modeled as a single node within the network, with the edges representing the inter-regional fibers of the brain. Consequently, the simulation comprises multiple nodes, each of which models the neural population activity in a specific brain region through a neural mass model. Additionally, each node includes several state variables governed by the intrinsic dynamics of the corresponding region. Therefore, if the simulated network consists of *m* nodes, each with *n*_*i*_ state variables, the total number of state variables would be 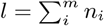. Furthermore, the edges of the network are defined using a connectivity matrix, commonly known as a connectome. The stochastic differential equation that represents the brain network model is expressed as follows:

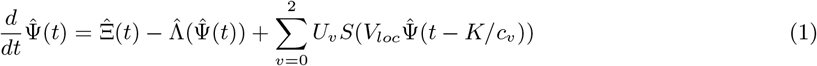

where 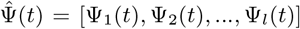 represents the vector of neural activity, with Ψ_*i*_(*t*) denoting the neural activity of the *i*^*th*^ node. The term 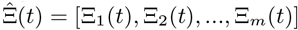 corresponds to vector of external inputs, such as noise or stimulus, with Ξ_*i*_(*t*) representing the external interventions for the *i*^*th*^ node. 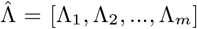 defines the local dynamics which also links the state variables within each neural mass. The last term in equation 1 denotes the coupling between state variables, where *U*_*v*_ represents the connectivity at different levels. Specifically, *U*_*v*=0_ refers to the local connectivity among the state variables within the same node, *U*_*v*=1_ represents short-range connectivity among nodes within the same region, and *U*_*v*=0_ corresponds to long-range connectivity among nodes across different regions, as defined by the connectome. In our region-based simulation, each region is represented by a single node, resulting in the absence of short-range connectivity (*U*_*v*=1_ = 0). Finally, *K* refers to the matrix of inter-regional distances, and *c*_*v*_ is the speed at which neural signals propagate through inter-regional fibers. The time delay for signal transmission between regions is accordingly calculated as *t*_*lag*_ = *K/c*_*v*_. However, for the local connectivity (*v* = 0), the speed is *c*_*v*_|_(*v*=0)_ = *∞*, implying that there is no time delay.

To start the simulation, the first step involves defining the connectivity matrix. At this stage, the region names, their coordinates, areas, tract/fiber lengths between each pair, and the directed structural connectivity matrix was determined. When defining the structural connectivity matrix, if the data is derived from anatomical tracing, the matrix is asymmetric which due to the fact that the connection strength from node *i* to node *j* differs from that from *j* to *i*. For diffusion-weighted imaging, however, the data lacks directionality, resulting in a symmetric connectivity matrix.

We simulated several scenarios containing networks of 6, 10 & 20 nodes, commonly referred to as regions-of-interest (ROIs). The simulated networks follow a particular structure, comprising two sub-networks: one activated by a task (task sub-network) and the other remaining at rest (rest sub-network). The connectivity pattern for each sub-network was designed to mimic a “small-world” topology. The connection strength was randomly assigned, ranging between 2 and 6, where each element (*i, j*) specifies the presence of a directed connection from node *j* to node *i*. Regarding the task signal, two identical external stimuli were applied to ROIs 1 & 2. Figure 1 (a.) illustrates graphical model of the simulated network, where each circle represents a region and the square indicates the stimulus, and the corresponding structural connectivity (Fig. 1 (b.)).

**Figure 1:**
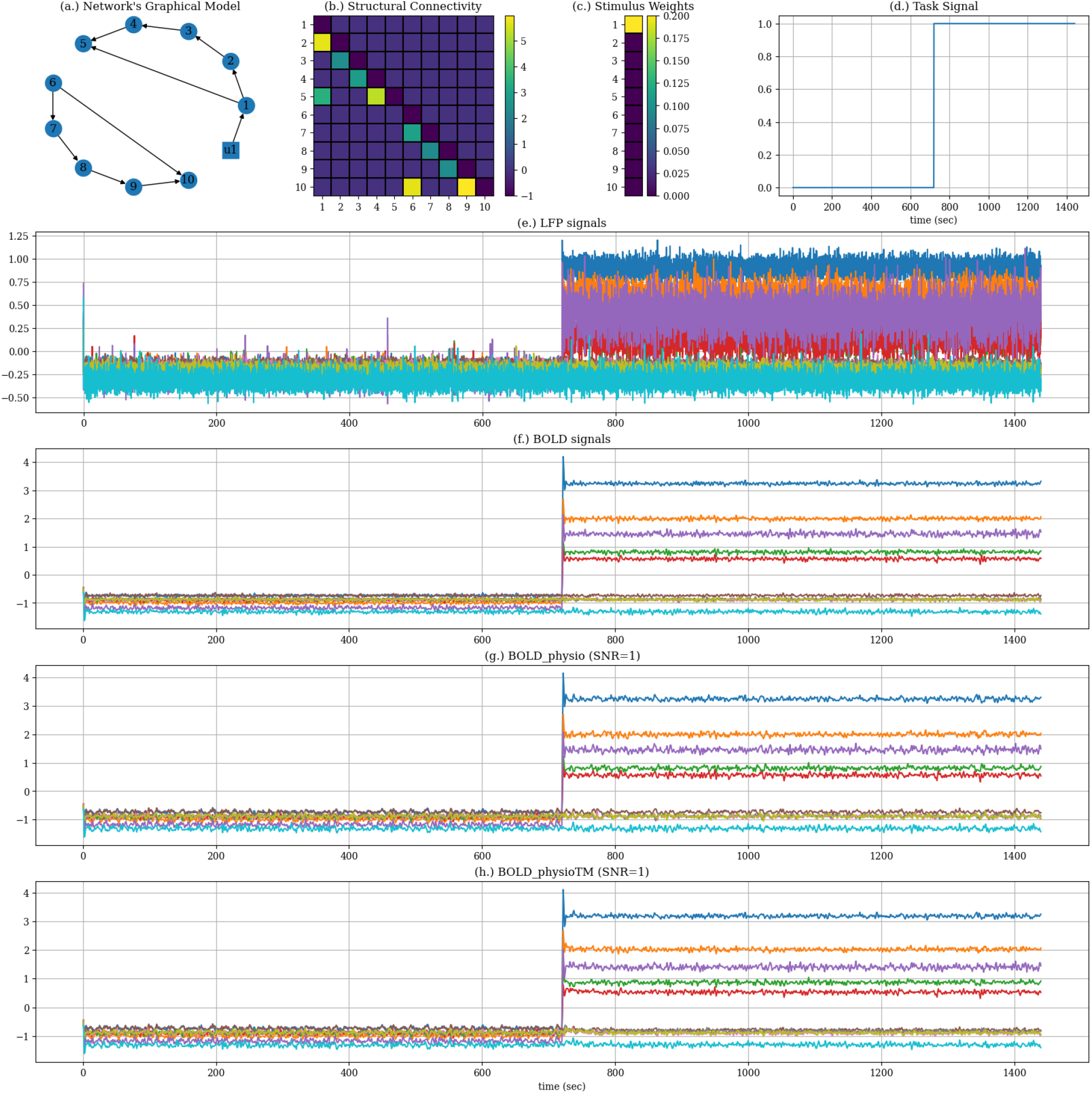
Simulation parameters and simulated Datasets: (a.) The graphical model of the simulated networks, (b.) the directed structural connectivity (SC) matrix, (c.) stimulus weights: the weights by which the stimulus was fed into each region, (d.) task/stimulus signal, (e.) simulated LFP signals, (f.) BOLD_raw: BOLD signals without physiological artifacts containing only neural signal, (g.) BOLD_physio: realistic BOLD signals with physiological artifacts, and (h.) BOLD_physioTM: BOLD signals with task-modulated physiological artifacts

Once the overall configuration of the network, including regions and connectomes, is established, we proceed to define the local dynamics that control the oscillation of state variables within each region. As supported by several studies [16, 17, 18], two-dimensional (2D) dynamical systems are capable of simulating various phenomena observed in neuronal populations, and many neural models can be formulated using 2D dynamical equations. Therefore, we employed the “*Generic*2*dOscillator*” formulation as described below;

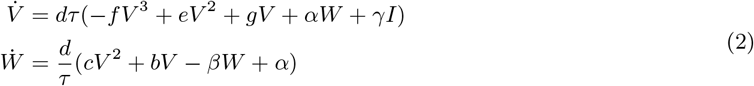

In this formulation, *V* and *W* denote the fast and slow oscillating state variables, respectively. The parameter *I* represents the baseline excitability of the system and serves as the entry point for external inputs, whether they are stimuli or signals from other regions. All other parameters are constants, set to values that produce dynamics similar to those described by the FitzHugh-Nagumo [16, 17].

In the following step, given that our focus is on a task-based problem, it is necessary to define an external stimulus. In this study, the task is specified at the regional level in both time and space. Spatially, Only the first region is affected by task. Accordingly, as shown in Figure 1 (c.), the stimulus weight is non-zero only for the first region. In time domain, the stimulus is modeled as a pulse train signal with an amplitude of 0.2, a period of 24 minutes, and a duty cycle of 50% (Fig.1 (d.)).

A Gaussian white noise with a standard deviation of 0.1% is also introduced as an internal noise to each region, representing their baseline neural activity. We employed Heun’s method, also known as the improved/modified Euler’s method, as the integrator to solve the equations forward in time, with a time step of 0.1 milliseconds. The simulated data is then down-sampled to a time step of 100 milliseconds which is referred to as our LFP data (Fig. 1 (e.)). The total simulation length is 24 minutes, beginning with a 12-minutes rest (task-off) period followed by a 12-minutes task activity (task-on period). With a time step of 100 milliseconds (*dt*_*LFP*_ = 0.1*sec*), this results in a total of 14, 400 data points.

### 2.2 Blood-Oxygenation-Level-Dependent (BOLD)

signal In the next step, we generated the blood-oxygenation-level-dependent (BOLD) signal using the corresponding simulated LFP signals. For this purpose, TVB utilizes a hemodynamic model, more specifically the Balloon model [19], to compute changes in blood flow and the concentration of oxy-/deoxy-hemoglobin in response to neural activity. In other words, the hemodynamic response function (HRF) serves as a link between neural activity and cerebral blood flow. The HRF used in this study can be expressed as follows:

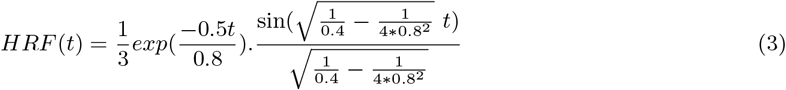

The simulation length for the BOLD data matches that of the LFP signals (24 minutes). With a repetition time (*T*_*r*_) equal to 1 minute, this resulted in the recording of 1, 440 BOLD data points, consisting of 720 rest and 720 task points (Fig. 1 (f.)). This simulated BOLD signal is hereafter referred to as the “*BOLD_raw*”, indicating that no noise or artifacts were added.

In the following section, we will discuss the physiological artifacts present in the BOLD signal and explain how we generated and integrated them into our “*BOLD_raw*”.

### 2.3 Physiological Artifacts

Physiological processes are known to induce systemic fluctuations, referred to as systemic low-frequency oscillations (SLOs), into the hemodynamic. To make the simulation more realistic, we generated physiological processes, including respiration and cardiac signals, and incorporated their artifacts into the simulated BOLD signals. The whole simulated signals are shown in Figure 2.

**Figure 2:**
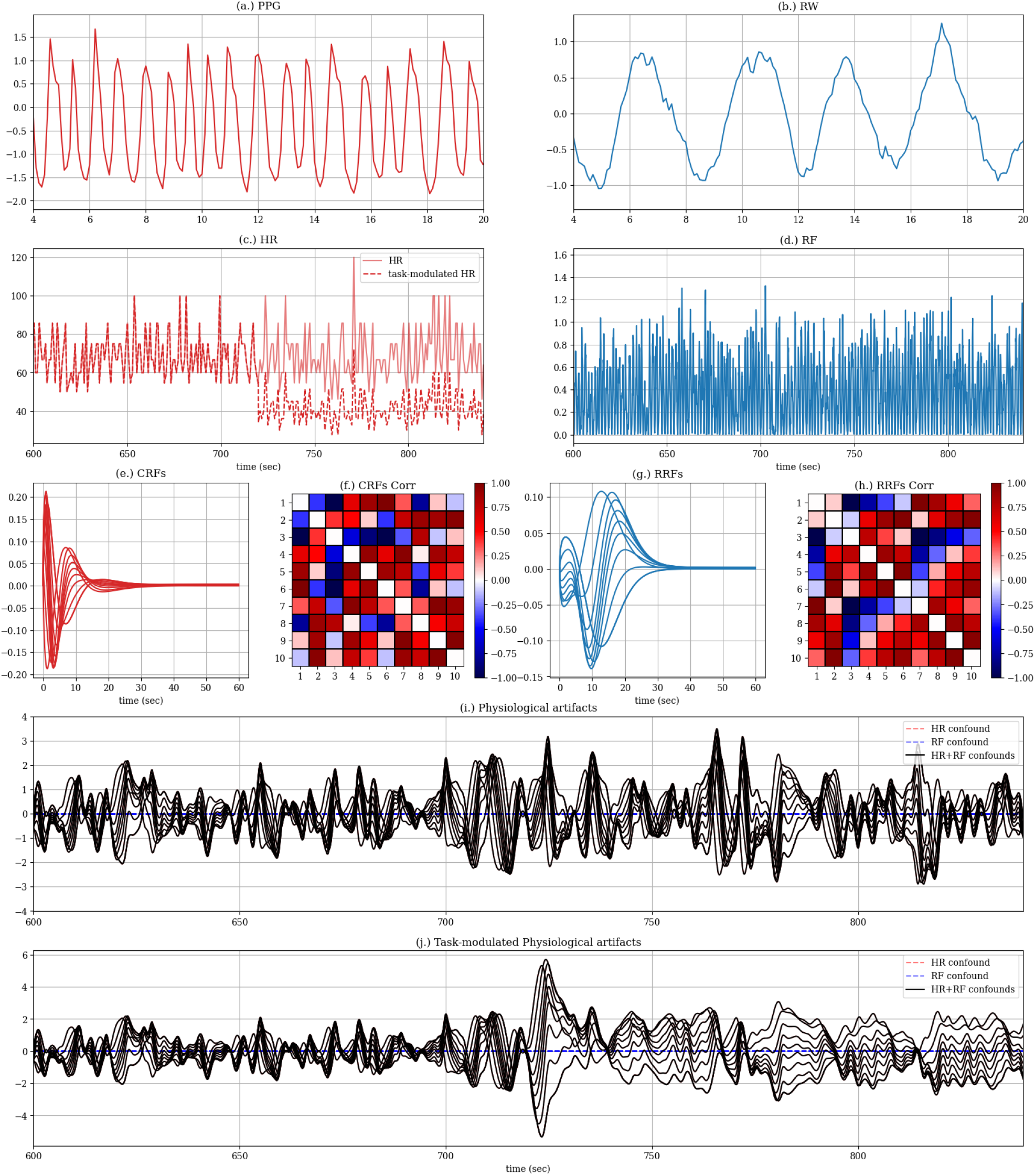
Physiological signals simulation: (a.) photoplethysmography (PPG), (b.) respiration wave-form (RW), (c.) heart rate (HR), (d.) respiration flow (RF), (e.) cardiac response functions (CRFs), (f.) correlation of CRFs, (g.) respiration response functions (RRFs), (h.) correlation of RRFs, (i.) resulting physiological artifacts; cardiact and respiration artifacts, (j.) task-modulated physiological artifacts

#### 2.3.1 Respiration and Cardiac Signals

We adopted the formulation proposed by Mann-Krzisnik and Mitsis (2022) [20]. The respiration waveform (RW) was generated using standard Stuart–Landau oscillators (SLOs). The differential equation governing the standard SLOs in Cartesian coordinates is given as follows;

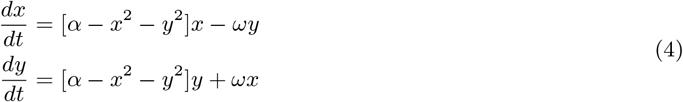

In this equation, *x* and *y* are two variables interacting through a nonlinear equation, with *x* representing the RW. The bifurcation parameter *α* is set to 0.75, and the natural angular frequency *ω* is set to 0.6*π*.

The cardiac signal, as observed in photoplethysmography (PPG) recordings, was simulated using the non-linear equation proposed by Rundo et al. (2018) [21]. To enhance the realism of the cardiac signal, the time constant for the PPG oscillator (Eq.5) was modulated by the respiration waveform (RW), calculated using Eq.4, to replicate the effect of respiratory sinus arrhythmia. This coupling between respiratory and cardiac activity leads to an increase in heart rate during inhalation and a decrease during exhalation in healthy adults [22].

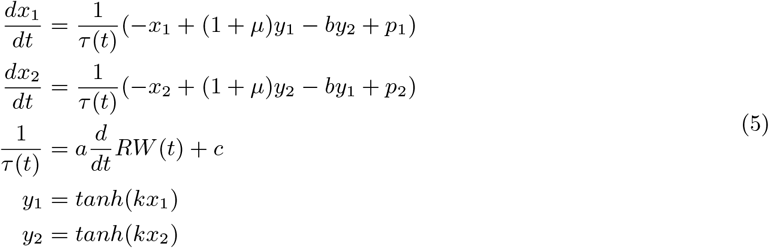

where *x*_2_ was designated as the PPG signal, and the time-constant *τ* (*t*) was directly modulated by the RW. The model parameters were set as follows: *μ* = 0.5, *b* = 1.0, *p*_1_ = −0.3, *p*_2_ = 0.3, *a* = 1.0, *c* = 14, and *k* = 1.5.

The PPG and RW signals were simulated using a stochastic Stratonovich–Heun algorithm [23], and are presented in Figure 2 (a.) and (b.), respectively. The heart rate (HR) signal was derived from the generated PPG by taking the reciprocal of the time interval between consecutive cardiac peaks, as shown in Figure 2 (c.). To calculate the respiratory flow (RF), the temporal derivatives of the RW were squared (Fig. 2 (d.)).

To translate the physiological signals into physiological artifacts in the BOLD signal, the generated signals must be convolved with physiological response functions (PRFs), as detailed in the following sections. Finally, the resulting artifacts were added to the BOLD signal at varying signal-to-noise ratios (SNRs) leading to a more realistic signal, hereafter referred as to “*BOLD_physio*”.

#### 2.3.2 Task-Modulated Physiological Signals

Given that we are examining task-based data, it is crucial to consider how stimulus impacts various parameters and analyze its effects on brain signals from different perspectives. Different task paradigms can influence BOLD signals in diverse ways; for instance, some tasks, such as visual tasks, primarily modulate neural signals, while others, like hand grip which increases the amplitude of the PPG signal (so-called physiological tasks), affect both neural signals and physiological processes. In the latter case, the physiological signals are also modulated by the task, which we refer to as “task-modulated physiological signals” in this study. To explore this scenario, we modulated the heart rate using the same task signal defined in TVB, simulating the impact of physiological tasks according to which the HF decreases by task initiation. The resulting task-modulated heart rate is shown in Fig. 2 (c.).

As detailed in section 2.3.1, the task-modulated physiological signals were integrated into the BOLD signals at different SNRs, after being convolution with the corresponding PRFs. The resulting signal, simulating what we would expect to record during physiological tasks, is hereafter referred to as “*BOLD_physioTM*.”

#### 2.3.3 Physiological Response Function

Physiological response functions (PRFs) were used as convolution models to convert the physiological processes into artifacts within the BOLD signal. Using the previously defined physiological signals, the resulting artifacts are as follows:

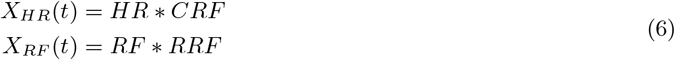

In Eq. 6, *CRF* and *RRF* refer to the cardiac and respiration response functions, respectively, while *X*_*HR*_ and *X*_*RF*_ represent the corresponding artifacts in the BOLD signal. Standard forms of *RRF* and *CRF* were introduced by [24] and [25], respectively. However, as suggested by Kassinopoulos & Mitsis (2022) [26], using a double gamma function can enhance performance compared to the standard models. The gamma function is defined as follows:

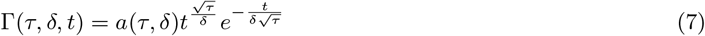

where, *τ* and *α* control the approximate time of peak and the dispersion of the function, while *α* is the scaling factor that normalizes the peak value of the gamma function to 1. Consequently, the PRF curves are defined as follows:

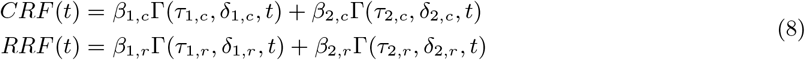

In total, equation 8 encompasses eight parameters for the gamma functions (*τ*_1,*c*_, *δ*_1,*c*_, *τ*_2,*c*_, *δ*_2,*c*_, *τ*_1,*r*_, *δ*_1,*r*_, *τ*_2,*r*_, *δ*_2,*r*_), and four scaling parameters (*β*_1,*c*_, *β*_2,*c*_, *β*_1,*r*_, *β*_2,*r*_). These parameters were randomly selected according to the recommendations of Kassinopoulos & Mitsis (2022) [26]. Accordingly, we generated the same number of CRFs and RRFs as the nodes in our networks, each uniquely assigned to a ROI (Fig. 2 (e.) & (g.)). Since this study investigates the relationships between regions, we also calculated the correlations between the PRFs to assess their influence as well. Figure 2 (f.) & (h.) illustrates the correlation of the generated PRFs.

Finally, by incorporating the generated physiological artifacts into the BOLD_raw signals, which reflects only the brain’s neural activity, we produced more realistic BOLD signals (BOLD_physio and BOLD_physioTM), as illustrated in Figure 1 (g.) & (h.).

## 3 Methods: Functional Connectivity (FC)

Functional connectivity (FC) is mathematically defined as temporal dependencies between spatially distinct brain regions. Numerous methods have been developed to assess FC, aiming to determine direct or indirect connections, as well as to explore linear or nonlinear relationships between two signals (*bivariate*) or among multiple signals (*multivariate*). FC can be analyzed in the time or frequency domain by evaluating the amplitude or phase values of the signals. External stimuli or tasks can modulate brain signals and their connections across different temporal scales, ranging from rapid neural responses (milliseconds) to minutes. Furthermore, these stimuli can also alter FC patterns that are typically observed during resting state at both neural [6] and system [7] level.

In this section, we will review the number of methods tested and provide a brief overview of each. It’s important to note that each method can be applied with different sets of controlling parameters. However, for this study, we have fine-tuned these parameters according to our data to obtain satisfactory results.

### 3.1 Correlation (Corr)

The Pearson correlation coefficient, referred to here as full correlation (Corr), is one of the most common and straightforward methods for assessing FC. This method evaluates the instantaneous linear relationship between two signals in the time domain based on their amplitudes values. In other words, correlation can be described as the covariance between two time series, normalized to have unit variance. The correlation coefficient between signals *x* and *y* is calculated as follows:

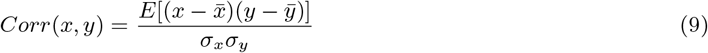

where *E*[*x*] represents the expected value of *x*, 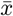 and 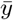 are the mean values of signals *x* and *y*, and *σ*_*x*_ and *σ*_*y*_ denote their standard deviations. The correlation results range from +1, indicating complete synchrony or correlation, to −1, signifying complete anti-correlation. A correlation value of zero between two signals implies no linear instantaneous relationship between them.

Full correlation is, however, a pairwise method which means it cannot differentiate between direct and indirect interactions or connections within a network of interconnected signals. To address this limitation, partial correlation methods have been developed.

### 3.2 Partial Correlation (Pcorr)

Partial correlation (PCorr) is defined as the correlation between two time series after removing the influence of all other time series in the dataset. This method offers a computationally efficient alternative to structural equation modeling (SEM, [27]). In our study, we applied an effective approach introduced by Marrelec et al., (2006) [28] to compute partial correlations of BOLD signals using the inverse covariance matrix, as shown below:

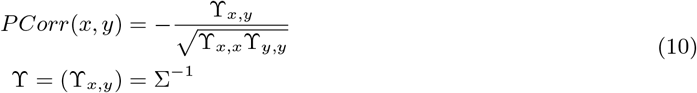

where Σ represents the covariance matrix calculated using all time series. The resulting values range between −1 and +1, similar to full correlation. However, a partial correlation value of zero between two regions indicates the absence of a direct connection between them.

### 3.3 Precision (Prec)

As previously mentioned, partial correlation can be effectively computed using the inverse of the covariance (ICOV) matrix, often referred to as the precision (Prec) matrix. As a network estimation method when applied to multivariate Gaussian data, the ICOV matrix is expected to be sparse. We achieved this sparsity by employing a regularization method such as Lasso ([29, 30]). By applying an *L*_1_ penalty to the estimated ICOV matrix, entries close to zero are shrunk more than others, thus satisfying the sparsity condition. This is especially beneficial when the number of observations per node is relatively small compared to the number of nodes in the network [13]. The mathematical formulation of the regularized ICOV, as presented by Friedman et al. (2008) [30], is formulated as follows:

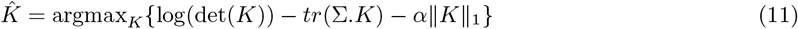

where *K* is the estimated precision matrix, Σ is the covariance matrix of the data. Moreover, *tr*(*·*) denotes the trace function (i.e., *tr*(*A*) =∑*A*_*ii*_), and |*K*|_1_ represents the *L*_1_-norm, which is the sum of the absolute values of the elements of matrix *K*. The regularization parameter *α* is set to 0.05 in this study.

### 3.4 Coherence (Coh)

Coherence (Coh) can be defined as the counterpart of correlation in the frequency domain. In other words, two signals are considered coherent if their power spectra are correlated. Coherence at a single frequency can be computed as follows:

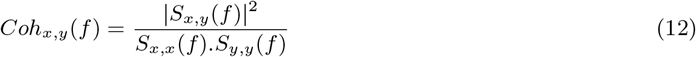

where *S*_*x,x*_ and *S*_*y,y*_ represent the power spectral densities of signals *x* and *y*, respectively, while *S*_*x,y*_ denotes the cross-spectral density of *x* and *y*, computed using Welch’s method [31]. For this study, we applied a *Hann* window of length 15 for Welch’s method. Coherence is typically calculated for a specific frequency or across multiple frequency bands, with the results subsequently combined. Smith et al. (2011) [13] found that averaging coherence across frequencies enhances network identification. However, due to differences in repetition time between our BOLD data (*TR* = 1 sec) and the data used by Smith et al. (*TR* = 3 sec), we limited our frequency band to *≈* 0.15*Hz*.

### 3.5 Mutual Information (MI)

Mutual information (MI) is a non-parametric, information-based method capable of capturing both linear and nonlinear statistical dependencies between two signals by quantifying the shared information between them. MI is grounded in entropy estimation from k-nearest neighbor distances [32] and is, in fact, closely related to Shannon entropy [33]. Due to its non-parametric nature, MI requires a larger number of samples for accurate estimation. For two continuous variables, MI can be expressed as:

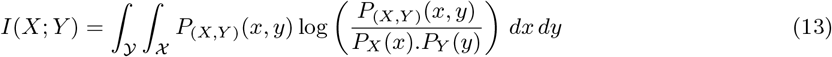

In this formula, *P*_(*X,Y*)_ represents the joint probability density function of *X* and *Y*, while *P*_*X*_ and *P*_*Y*_ denote the marginal probability density functions of *X* and *Y*, respectively.

### 3.6 Zero-lag Regression (0-lagReg)

Zero-lag (instantaneous) regression is a semipartial, multivariate method based on general linear models (GLM). This approach estimates the level of effective (direct) connectivity between each pair of regions while accounting for the effects of other ROIs. In some studies, this method is also referred to as multivariate ROI-to-ROI connectivity (mRRC) [35, 36]. Specifically, this method focuses on assessing connectivity at the level of ROIs by modeling direct influences between them, thereby offering a more detailed view on the network structure than bivariate approaches. Zero-lag regression operates in the time domain and computes instantaneous connectivity using the amplitude values of signals. The mathematical formulation of zero-lag regression is given by:

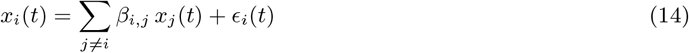

where *x* represents the time series extracted from different regions, and *β*_*i,j*_ is the multivariate regression coefficient that quantifies the instantaneous effect of region *j* on region *i*. These regression coefficients are then used as connectivity measures.

### 3.7 Granger Causality (GC)

Granger causality (GC), first introduced by Granger (1969) [37], is a directed/causal connectivity measure. The core idea behind this causality measure is based on the principle of temporal precedence, which states that “a cause must precede its effect.” This concept allows us to identify the direction of interactions among different variables of a system. Practically speaking, variable *X* is said to “Granger cause” variable *Y* if knowing the past values of *X* improves the prediction of *Y* beyond what is possible by only knowing the past values of *Y*. Granger causality is typically implemented using autoregressive (AR) models, where the time series data are modeled as a function of their own past values and the past values of other variables in the system. In this study, we explored three different variants of Granger causality: Geweke’s conditional GC, directed transfer function (DTF), and pairwise GC. Each of these methods is explained in the following sections.

#### 3.7.1 Pairwise GC (pwGC)

In 1982, John Geweke [38] adapted the concept of Granger causality, originally utilized to analyze relationships between two stochastic processes, to be applicable within linear autoregressive models. Consider *X* and *Y* as two stationary time series, where their mean and variance do not depend on time (*t*). The time series *x*(*t*) and *y*(*t*) can be estimated using their own past values as follows:

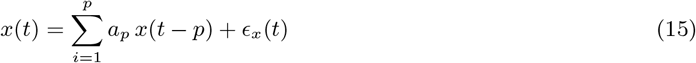

and

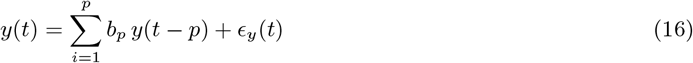

where *a* and *b* represent the coefficients of the model, while *ϵ*_*x*_ and *ϵ*_*y*_ are the residuals (model prediction errors) for each time series. The parameter *p* is the maximum number of lagged observations included in the model, which is determined using the Akaike Information Criterion (AIC, [39]) and Bayesian Information Criterion (BIC, [40]) as follows:

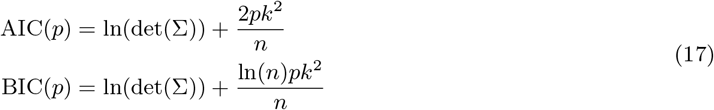

In Eq. 17, Σ is the covariance matrix of the residuals for the corresponding order-*p* model, *p* is the number of lagged observations, *k* is the number of variables included in the model, and *n* is the number of time-points (observations). The model order is selected based on the point where the AIC or BIC is minimized. In this study, we used the BIC to select the model order. Furthermore, using their pair-wise autoregressive models, *x*(*t*) and *y*(*t*) can be estimated as follows;

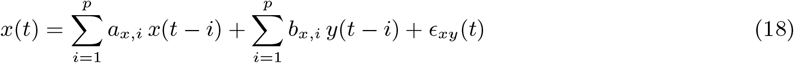

and

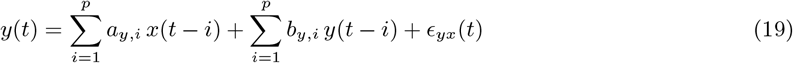

Since there is no zero-lag term in Eqs. 18 & 19, and the variables do not exhibit instantaneous interactions, this model is referred to as a “simple causal model.” If a model includes zero-lag terms, it is known as a model with instantaneous causality. Referring to the fundamental concept of Granger causality, the measure of linear feedback (causality) from *X* to *Y* and vice versa can be defined as follows:

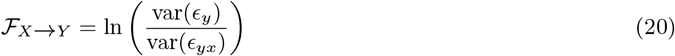

and

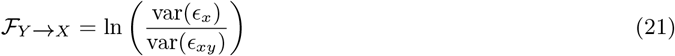

This measure, which determines the causal interactions between two time series, is considered a bivariate method. However, to account for the influence of other variables, Geweke introduced an extended method (1984) [41].

#### 3.7.2 Geweke’s Conditional GC (GGC)

Following his earlier work, Geweke introduced another measure in 1984 to compute the conditional linear dependency between time series [41]. Consider a multivariate time series *X* = [*x*_1_, *x*_2_, *x*_3_] with three variables/channels and *n* observations/samples. The full, unrestricted multivariate autoregressive (MVAR) model, similar to the one described in Section 3.7.1, that includes all the variables in *X*, can be defined as follows:

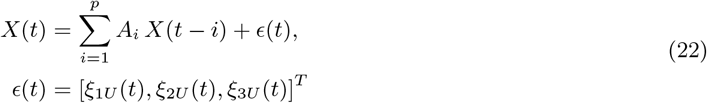

where *A*_*U*_ is the coefficient matrix for the unrestricted model, *p* is the model order determined by BIC, and *ϵ*(*t*) represents the vector of residuals at time *t*. To determine the conditional feedback from variable 2 to variable 1, the restricted model is defined by excluding the time series of variable 2 as follows:

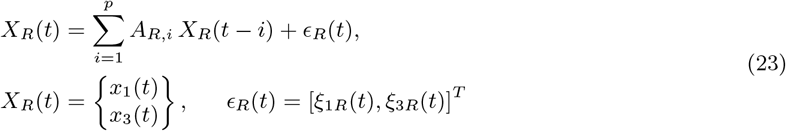

Finally, the Granger causality from variable 2 to variable 1, conditioned on variable 3, is given by;

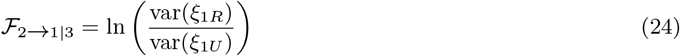

This method can identify directed connections among nodes in a network using time-domain amplitude values. Additionally, this approach is scalable and can be extended to systems with a higher number of variables.

#### 3.7.3 Directed Transfer Function (DTF)

As another approach to obtaining Granger causality, the Directed Transfer Function (DTF) uses the coefficients of the MVAR model to study GC. Initially introduced by Kami’nski [42], DTF describes the propagation of brain signals between EEG channels by revealing their directional flow and frequency content. We begin by reformulating the MVAR model of Eq. 22 as follows:

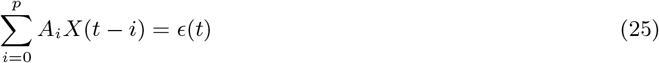

where *A*_*i*_ represents the matrix of coefficient estimates for lag *i*. It is important to note that to satisfy the “simple causal model” condition (i.e., no instantaneous causality), *A*_0_ should be equal to the identity matrix (*I*). The model order (*p*) is computed using BIC. Next, to estimate the spectral properties of the processes under study, the signals are transformed to the frequency domain using the *Z*-transform as follows:

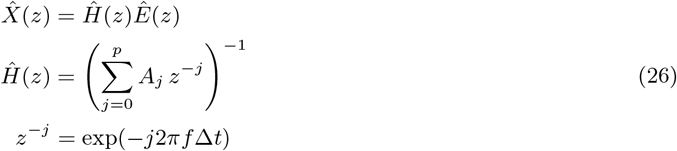

where *Ĥ* (*z*) is the transfer function of the system and *z*^−1^ represents a unit delay operator. The matrix *Ĥ* is not symmetric, meaning that the transition from channel *i* to *j* is different from the transition from channel *j* to *i*. This asymmetry allows the study of directionality using this method. Finally, to enable comparison across different channels, the transfer function i s normalized as follows:

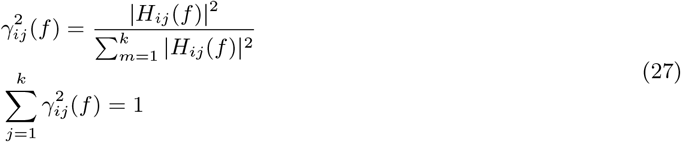

Here, *γ*_*i,j*_ (*f*), the directed transfer function, is a normalized measure of directional connectivity among different variables as a function of frequency (*f*). DTF is closely related to coherence. Similarly to coherence, we averaged the DTF results over frequencies up to *≈* 0.15*Hz* to obtain a single FC matrix.

## 4 Results & Discussion

This study centers on two principal objectives. First, we assessed the capacity of different Functional Connectivity (FC) methods to detect various types of connections within a network, using Local Field Potential (LFP) and Blood-Oxygen-Level-Dependent (BOLD) signals. TO begin with, we defined three types of connections between each pair of regions: direct connections (DC), indirect connections (iDC), and functional connections (FnC). A direct connection (DC) refers to a neural pathway with an uninterrupted anatomical link between two brain regions. Within the context of the network under study, a direct connection is classified as a true positive based on the structural connectivity matrix (Fig. 3 (c.)). An indirect connection (iDC) refers to a pathway where the connection between two brain regions is mediated through one or more intermediate regions (Fig. 3 (d.)). Finally, a functional connection (FnC) is defined as any connection that links two regions, either directly or indirectly, enabling them to function together. In other words, if there is a pathway from one region to another, a functional connection exists between them (Fig. 3(e.)). Based on these definitions, if there is neither direct nor indirect pathway between regions, the pair is considered to have no connection (NC) (see Fig. 3 (f.)). Consequently, for the network under study, any connections among the regions across different sub-networks are regarded as NC. Given that only some of the methods employed can estimate directionality (resulting in asymmetric matrices), we restricted our analysis to the lower triangle of the FC matrices. Figure 3 provides a summary of these definitions.

**Figure 3:**
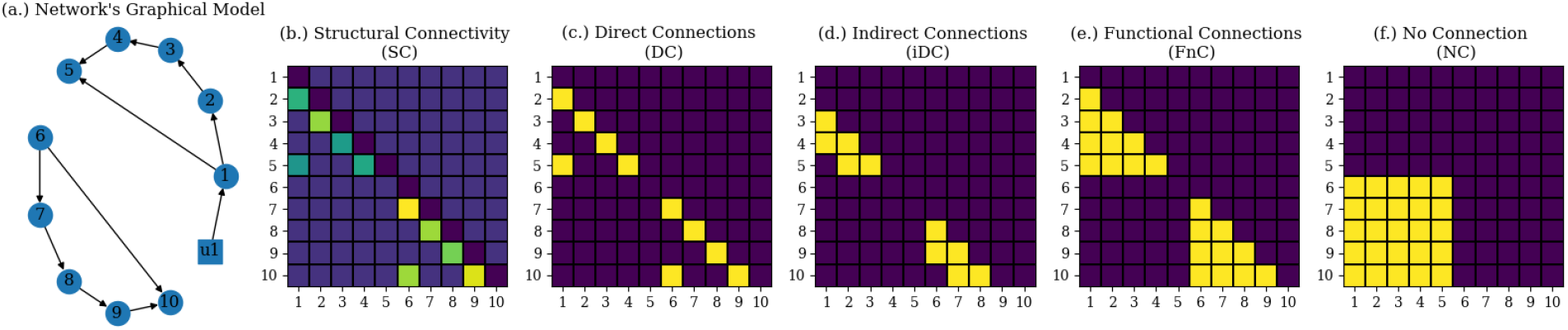
Different types of connections in the network under study; (a.) the graphical model of the network, (b.) structural connectivity matrix, (c.) direct connections (DC), (d.) indirect connections (iDC), (e.) functional connections (FnC), (f.) no connection (NC)

Secondly, we introduced physiological artifacts (as explain in Section 2.3) to evaluate the reliability and robustness of these methods under different conditions, including various task paradigms, and varying noise levels. We simulated various scenarios for this study, comprising different task paradigms, noise levels, network sizes (varying numbers of nodes), and the application of either unique PRFs for each region or a identical PRF across all regions.

It is important to note that for each scenario, we simulated 20 subjects with the same network structure but varying structural connectivity values (ranging from 2 to 6). All simulated BOLD signals consist of three datasets: “BOLD_raw” which contains only neural signals, “BOLD_physio” which contains neural signals and the physiological artifacts, and “BOLD_physioTM” which contains neural signals and the task-modulated physiological artifacts. In the latter two datasets, various SNRs are applied, as outlined in Table 1. A summary of these simulation scenarios is provided in Table 1.

**Table 1:** Summary of the simulations

| Sim. # | Signal modality | # nodes | Other factors |
| --- | --- | --- | --- |
| Sim. 1 | LFP | 10 | - - - |
| Sim. 2 | LFP | 6 | - - - |
| Sim. 3 | LFP | 20 | - - - |
| Sim. 4 | BOLD_raw | 10 | - - - |
| Sim. 5-7 | BOLD_physio | 10 | Different PRFs, SNRs=[1,5,10] |
| Sim. 8-10 | BOLD_physioTM | 10 | Different PRFs, SNRs=[1,5,10] |
| Sim. 10-12 | BOLD_physio | 10 | same PRFs, SNRs=[1,5,10] |
| Sim. 13-15 | BOLD_physioTM | 10 | same PRFs, SNRs=[1,5,10] |
| Sim. 17 | BOLD_raw | 6 | - - - |
| Sim. 18-20 | BOLD_physio | 6 | Different PRFs, SNRs=[1,5,10] |
| Sim. 21-23 | BOLD_physioTM | 6 | Different PRFs, SNRs=[1,5,10] |
| Sim. 24 | BOLD_raw | 20 | - - - |
| Sim. 25-27 | BOLD_physio | 20 | Different PRFs, SNRs=[1,5,10] |
| Sim. 28-30 | BOLD_physioTM | 20 | Different PRFs, SNRs=[1,5,10] |

### 4.1 LFP-FC results: Method Comparison

Firstly, to mitigate the transient effect presents at the onset of each sequence, which synchronizes all signals, we excluded the first five seconds of data from the dataset. The remaining signals were then used to compute the FC matrices. Figure 4 (a.) presents a summary of the computed FC matrices using LFP signals. The structural connectivity matrix and the graphical model of the corresponding network (as described in Section 2) are displayed in Figure 3 (a.) & (b.). The FC matrices computed during the rest period are shown in the upper row, while those computed using task data are presented in the lower row.

**Figure 4:**
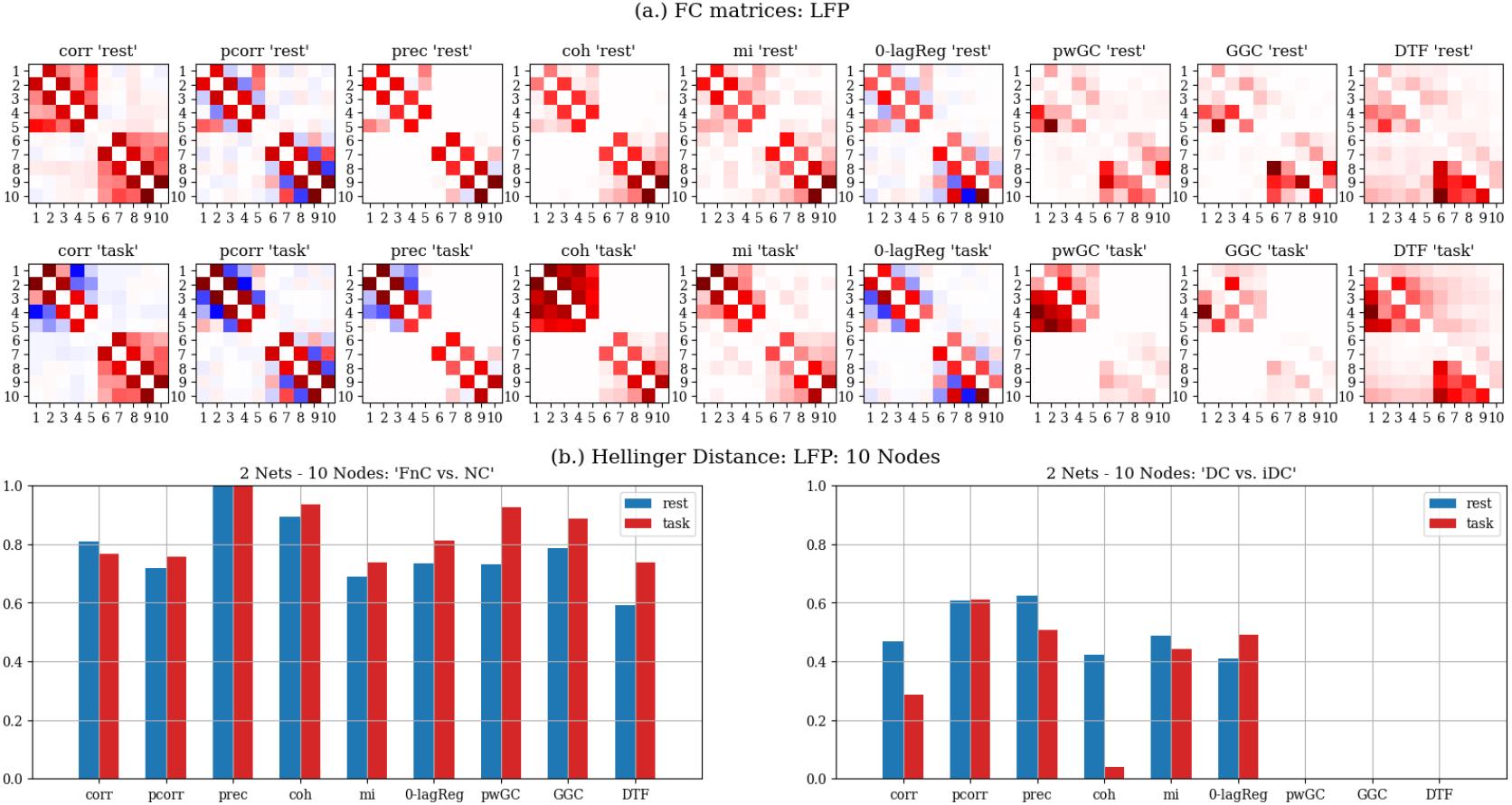
(a.) Rest- & Task-FC matrices computed using LFP signals, (b.) Hellinger distance: FnC vs NC, and DC vs iDC

Since each FC assessment method is based on distinct assumptions and calculated using different mathematical formulations, the resulting connection strengths have different scales. To improve the interpretability of these results, we compare the ability of each method to differentiate between various types of connections. For this purpose, we categorized the FC values into four groups based on their connection types and calculated the distribution of values for each category. To assess how well each method distinguishes between FnC and NC, as well as DC and iDC, we computed the “Hellinger” distance between the respective distributions. This distance quantifies the dissimilarity in the distributions of the FC values, with larger Hellinger distances indicating a greater ability to distinguish between connection types. Figure 4 (b.) presents the Hellinger distance values between FnC vs. NC and DC vs. iDC for each method during rest and task.

Starting with the distinction between FnC and NC, Figure 4 (b.) demonstrates that, in general, all methods performed acceptably, with precision (*prec*) showing the best performance during both rest and task periods. Moreover, except correlation, a general trend for all method is that all methods perform better during task compared to rest. Regarding the differentiation between DC and iDC, none of the *GC*-based measures were able to distinguish between these types of connection effectively. Among the remaining measures, partial correlation (*pcorr*) and *prec* are performing better than other methods methods during rest. During task, while the performance of some methods such as correlation (*corr*) and coherence (*coh*) drops, *pcorr* can still reliably distinguish the direct vs indirect connections. Overall, *pcorr* achieved the best dinstinction between DC and iDC.

### 4.2 BOLD-FC results: Method Comparison

We conducted a similar analysis on the BOLD data. The FC matrices computed using BOLD_raw dataset are displayed in Figure 5 (a.). When comparing these results with those obtained from LFP signals (Figure 4 (a.)), we observe a general increase in connectivity patterns across all measures in the BOLD dataset. Notably, BOLD-FC results, especially those derived from GC-based methods, exhibit higher inflation for across-network connections. This is unexpected, as these connections are not supposed to occur based on the structural connectivity matrix. This discrepancy can be attributed to several factors: the difference in temporal resolution between LFP signals (*dt* = 0.1*seconds*) and BOLD signals (*T*_*r*_ = 1*second*); and the fact that BOLD signals are generated by convolving LFP signals with the Hemodynamic Response Function (HRF), which acts as a low-pass filter. Consequently, the HRF and reduced temporal resolution smooth out rapid dynamics, leading to more homogeneous FC patterns.

**Figure 5:**
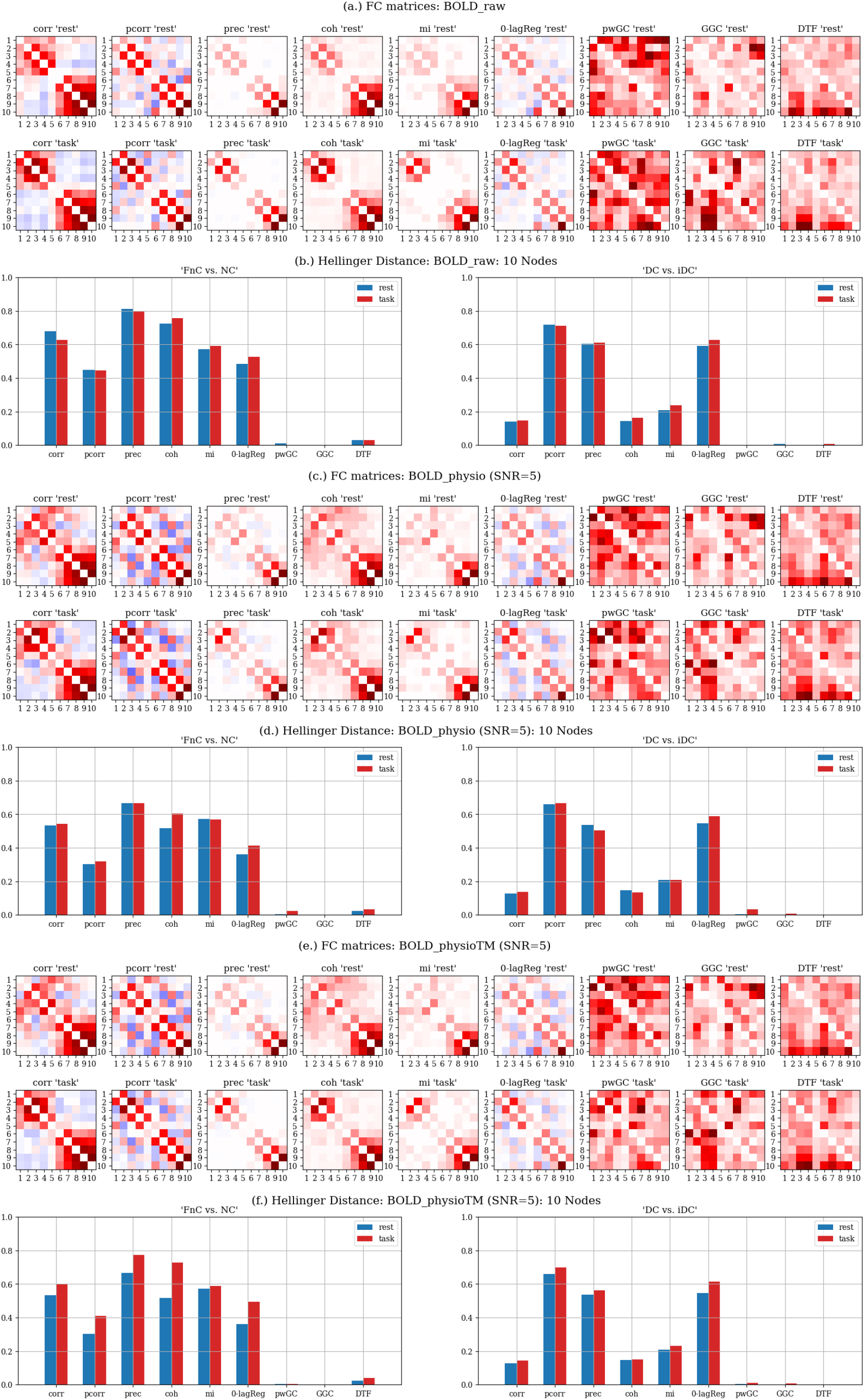
Rest- & Task-FC matrices computed using BOLD_raw (a.), BOLD_physio (c.), and BOLD_physioTM (e.) signals, and the Hellinger distance: between FnC vs NC, and DC vs iDC computed using BOLD_raw (b.), BOLD_physio (d.), and BOLD_physioTM (f.) datasets

Similarly to the approach used for LFP results (see subsection 4.1), we categorized BOLD-FC values into four groups (DC, iDC, FnC, and NC) and computed their distribution. To compare the distance between the corresponding distributions, we calculated Hellinger distance, shown in Figure 5 (b.).

As shown in Figure 5 (b.), none of the GC-based measures effectively distinguish between different types of connections, indicating that they may not be well-suited for analyzing BOLD data. This finding is consistent with the results reported by Smith et al. (2011) [13]. This limitation is likely due to the lower temporal resolution of BOLD data, which does not capture rapid temporal dynamics. In differentiating between FnC and NC, *prec* performs the best during both rest and task periods. However, *pcorr* is more effective at distinguishing between DC and iDC in both rest and task conditions. Additionally, an increase in the distance between connection types is observed during task performance, which can be attributed to the increased neural activity during this period.

### 4.3 Effect of Physiological Artifacts on BOLD-FC

In this section, we quantify the impact of physiological artifacts on FC results derived from realistic BOLD (BOLD_physio & BOLD_physioTM) signals across varying SNRs. Our analysis focuses on two primary task paradigms: physiological and non-physiological tasks. In the latter scenario, external stimulus exclusively modulate neural activity. Conversely, in physiological tasks such as hand grip, both neural signals and physiological artifacts are influenced by the task. The details of the simulation process are provided in subsection 2.3. In order to improve the interpretability of our results, we excluded respiratory artifacts from the BOLD signals, concentrating only on the effects of cardiac artifacts. Comprehensive results that include both types of artifacts are available in the supplementary materials.

Following the methodology outlined in previous sections, we computed FC measures using the BOLD_physio and BOLD_physioTM datasets, as shown in Figures 5 (c.) and 5 (e.), respectively. These results generally reveal a uniform increase in all FC patterns compared to those obtained from BOLD_raw data (Figure 5 (a.)). This increase is primarily attributable to the presence of physiological artifacts, specifically due to the synchronization of low-frequency oscillations in the BOLD signal. Notably, this effect is more pronounced in bivariate methods, where the influence of physiological artifacts is not compensated by regressing out other signals.

The computed FC values are categorized into four groups based on their connection types. The Hellinger distances between the distributions of FnC vs. NC and DC vs. iDC are then calculated. The results are presented in Figures 5 (d.) and 5 (f.).

Physiological artifacts generally reduce the ability of all methods to distinguish between different connection types. In both datasets (BOLD_physio and BOLD_physioTM), *prec* demonstrated the best performance in distinguishing FnC from NC during both rest and task, while *pcorr* performed better in distinguishing direct versus indirect connections. When comparing different task paradigms —physiological versus non-physiological— since the rest sequences is identical, it shows no differences, as expected. However, when comparing the task sequences between BOLD_physio and BOLD_physioTM, since the physiological artifact is modulated by the task in BOLD_physioTM (Fig. 5 (f.)), the variance of artifacts decreases with task initiation. This reduced variance during task significantly improves the distinction between FnC and NC for all methods, as compared to the same results with BOLD_physio dataset (Fig. 5 (d.)). However, for DC versus iDC, the methods’ performance does not improve significantly during task compared to rest. *pcorr* continues to show the best performance, likely due to its ability to compensate for physiological artifacts by regressing out other signals when computed between two regions.

### 4.4 BOLD-FC results with an Identical PRF across Regions

In this section, we present the results of a special case where physiological signals are convolved with the same CRF and then added to the neural BOLD signals from different regions. Figures 6 (a.) & 6 (c.) illustrate the FC results for both the rest and task periods.

**Figure 6:**
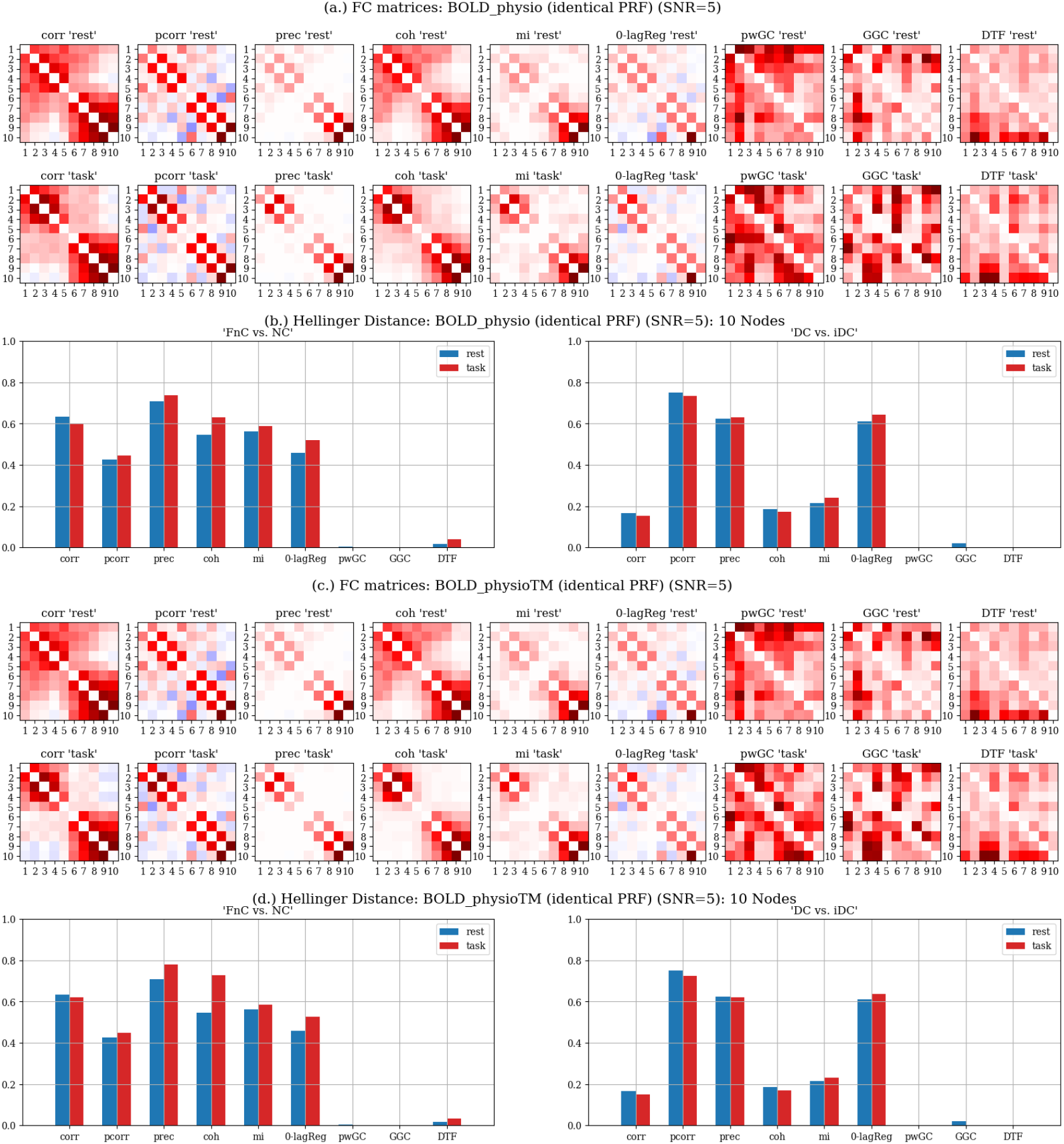
Rest- & Task-FC matrices computed using BOLD_physio (a.) and BOLD_physioTM (b.) signals with an identical PRF, and the Hellinger distance: between FnC vs NC, and DC vs iDC computed using BOLD_physio (c.) and BOLD_physioTM (d.) datasets

Following the same procedure used in the previous sections, the FC values were categorized into four groups considering their connection types (DC, iDC, FnC, and NC). The Hellinger distances between FnC versus NC, and DC versus iDC were computed and are presented in Figures 6 (b.) & 6 (d.).

Using a single PRF synchronizes the temporal variations of the added artifacts across different regions, effectively aligning the physiological artifacts temporally. This synchronization is expected to artificially inflate connectivity in FC values, particularly when using bivariate methods. In contrast, multivariate methods can compensate for these temporally aligned artifacts when regressing out signals from other regions. This phenomenon can be observed by comparing Figures 5 (b.), 6 (b.), and 6 (d.). The performance of *pcorr* in distinguishing FnC from NC, as well as DC from iDC, remains consistent across the BOLD_raw, BOLD_physio, and BOLD_physioTM datasets, even when physiological artifacts are temporally aligned, i.e., convolved with the same PRF for all regions. Additionally, by comparing Figures 5 (d.) and 6 (b.), it is evident that *pcorr* demonstrates better distinction in the identical PRF scenario. Besides, *prec* remains the best method for distinguishing FnC vs NC and a similar improvement can be observed.

### 4.5 FC Errors Induced by Physiological Artifacts vs. PRF Correlation

In this section, we investigate how the similarity of PRFs influences the magnitude of physiologically induced FC errors during both rest and task conditions across various methods. The FC matrices derived from the BOLD_raw dataset serve as the ground truth, representing a condition with no artifacts. To quantify the FC errors, we subtracted the connectivity matrices computed using the noisy BOLD datasets (BOLD_physio & BOLD_physioTM) from those obtained using BOLD_raw. The resulting FC errors between pairs of regions are plotted against the correlation of their respective PRFs. For a clearer visualization, we present results of four methods: *corr, pcorr, mi*, and *GGC*, as shown in Figure 7. The corresponding results for other methods can be found in the appendix. Additionally, a first-order regression line was fitted to quantify the relationship between FC errors and the CRF correlations, and is also presented in Figure 7.

**Figure 7:**
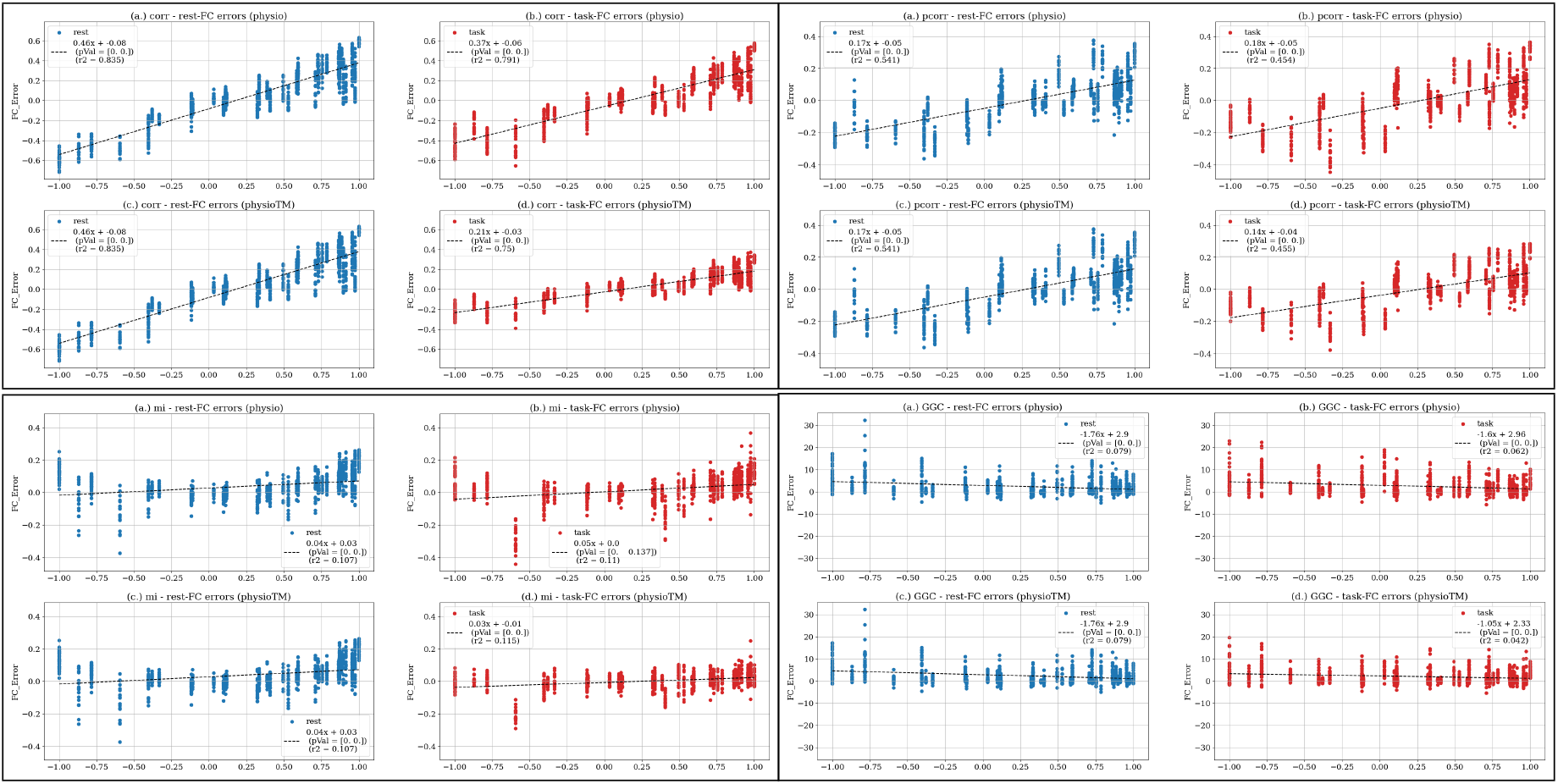
Errors induced by physiological artifacts to rest- & task-FC values vs. correlation of CRFs: over all BOLD simulations

Generally, task-FC error rates are equal to or smaller than rest-FC errors, particularly for methods that rely on temporal amplitude values of the signal which can be attributed to increased neural activity during task execution. Additionally, when comparing task-FC results from the BOLD_physio and BOLD_physioTM datasets, the latter —containing task-modulated physiological artifacts— exhibits reduced errors. This reduction is likely due to the decreased variance of physiological artifacts during the task condition. Furthermore, multivariate methods typically yield lower FC errors than bivariate methods during both task and rest periods, likely because they can effectively mitigate physiological artifacts by regressing out signals from other regions.

### 4.6 FC Errors Induced by Physiological Artifacts across Different SNRs

In this section, we examine how the SNR —the ratio of neural signals to physiological artifacts in BOLD data— affects rest- and task-FC values across different methods. For this analysis, We utilized two simulated datasets, BOLD_physio and BOLD_physioTM. As previously described, the FC errors that are induced by physiological artifacts were determined by subtracting FC matrices computed using noisy BOLD signals from those computed using BOLD_raw. Figure 8 presents the mean and standard deviation of the errors across varying SNR levels for *corr, pcorr, mi*, and *GGC*. Complete presentation of the results for all methods are provided in the appendix.

**Figure 8:**
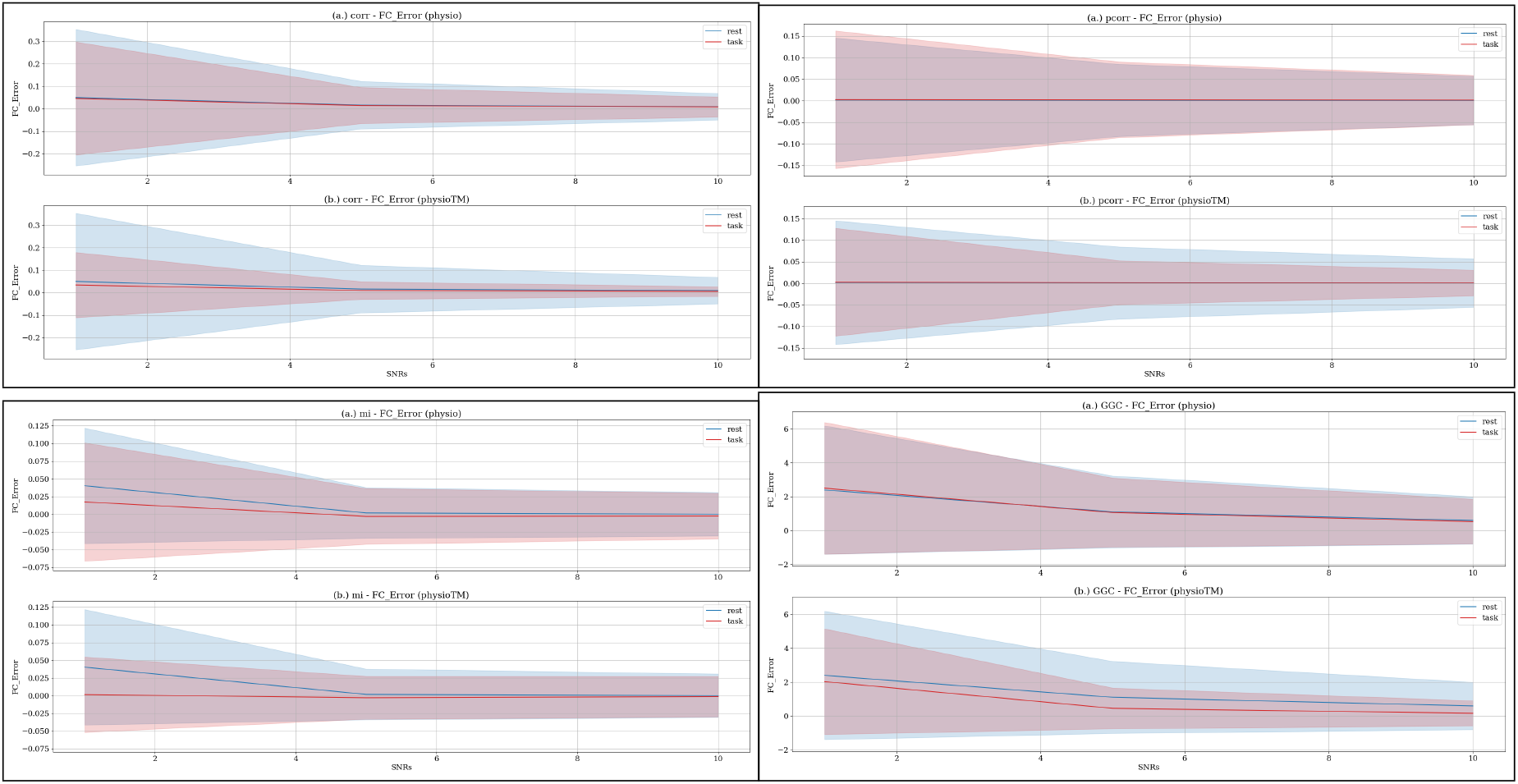
Mean and standard deviation band of the errors induced by physiological artifacts to rest- & task-FC values across various SNRs: over all BOLD simulations

As illustrated in Figure 8, higher SNR levels consistently lead to a reduction in both the mean and standard deviation of FC errors across all methods, as expected. When comparing *corr* with *pcorr*, the latter, being partially considered as a multivariate method, more effectively mitigates errors between pairs of regions by regressing out signals from other regions. Additionally, task-FC errors are generally equal to or lower than rest-FC errors at comparable SNR levels, likely due to increased neural activity during task performance. Furthermore, task-FC errors computed using the BOLD_physioTM dataset are significantly lower than those of the BOLD_physio dataset, which can be attributed to task-related modulation of physiological artifacts, reducing the impact of physiological noise in the BOLD signal.

## 5 Conclusion

This study provides a thorough evaluation of various Functional Connectivity (FC) methods, focusing on their performance with Local Field Potential (LFP) and Blood-Oxygen-Level-Dependent (BOLD) signals while accounting for physiological artifacts. Our analysis underscores the advantages of different FC measures and highlights critical considerations for enhancing connectivity measurements in neuroscience research.

Our investigation reveals that partial correlation is particularly robust in differentiating between direct and indirect connections in LFP datasets. Its ability to isolate direct connections by controlling for indirect pathways provides valuable insights into the structural and functional organization of neural networks. Precision and coherence also demonstrate strong performance in identifying functional connections, reflecting their capacity to capture dynamic interactions within the brain. Overall, all methods under study perform acceptably well in identifying functional connections, especially during task execution.

In contrast, the use of BOLD signals introduces unique challenges due to their lower temporal resolution and the inherent smoothing effect of the Hemodynamic Response Function (HRF). Our results indicate that Granger Causality (GC)-based methods face significant limitations in detecting any connectivity patterns under these conditions. This emphasizes the importance of carefully considering signal characteristics when interpreting FC results, and suggests that alternative methods, such as partial correlation, may provide more reliable connectivity estimates in the context of BOLD data.

The impact of physiological artifacts on FC measurements is a central focus of our study. These artifacts significantly distort the accuracy of connectivity estimates by inflating measurements and masking true neural interactions. Our analysis reveals that the effects are particularly detrimental when not properly regressed out, often leading to an overestimation of connectivity. However, with task-modulated physiological artifacts, some of these adverse effects are mitigated. By aligning physiological signals with task-related activity, the variance introduced by the artifacts is reduced, leading to improved performance of certain FC measures during the task period.

The study also demonstrates that FC values in regions with similar physiological response functions are more heavily impacted by physiological artifacts. This effect is particularly evident in the special case analysis where identical Physiological Response Functions (PRFs) were applied uniformly across brain regions. Using a single PRF across regions led to inflated connectivity measures due to the temporal alignment of physiological artifacts. However, multivariate methods showed resilience to these artifacts by effectively regressing out signals from other regions, maintaining consistent performance across different datasets. Additionally, our findings emphasize the importance of higher Signal-to-Noise Ratio levels in minimizing FC errors. Higher SNR consistently results in lower errors induced by physiological artifacts in connectivity measures, compared to those computed from the raw dataset, underscoring the critical need to optimize signal quality for accurate results.

Figure 9 summarizes the Hellinger distances between FnC vs NC, and DC vs iDC during rest and task periods across all 30 simulations. These simulations encompass varying network sizes (*m* = 6, 10, 20), different signal modalities (LFP and BOLD signals), diverse task paradigms with distinct impacts on physiological artifacts, and multiple noise levels. Precision consistently emerges as the best method for distinguishing functional connections in both rest and task conditions. On the other hand, partial correlation demonstrates the best performance in differentiating direct from indirect connections during both rest and task periods.

**Figure 9:**
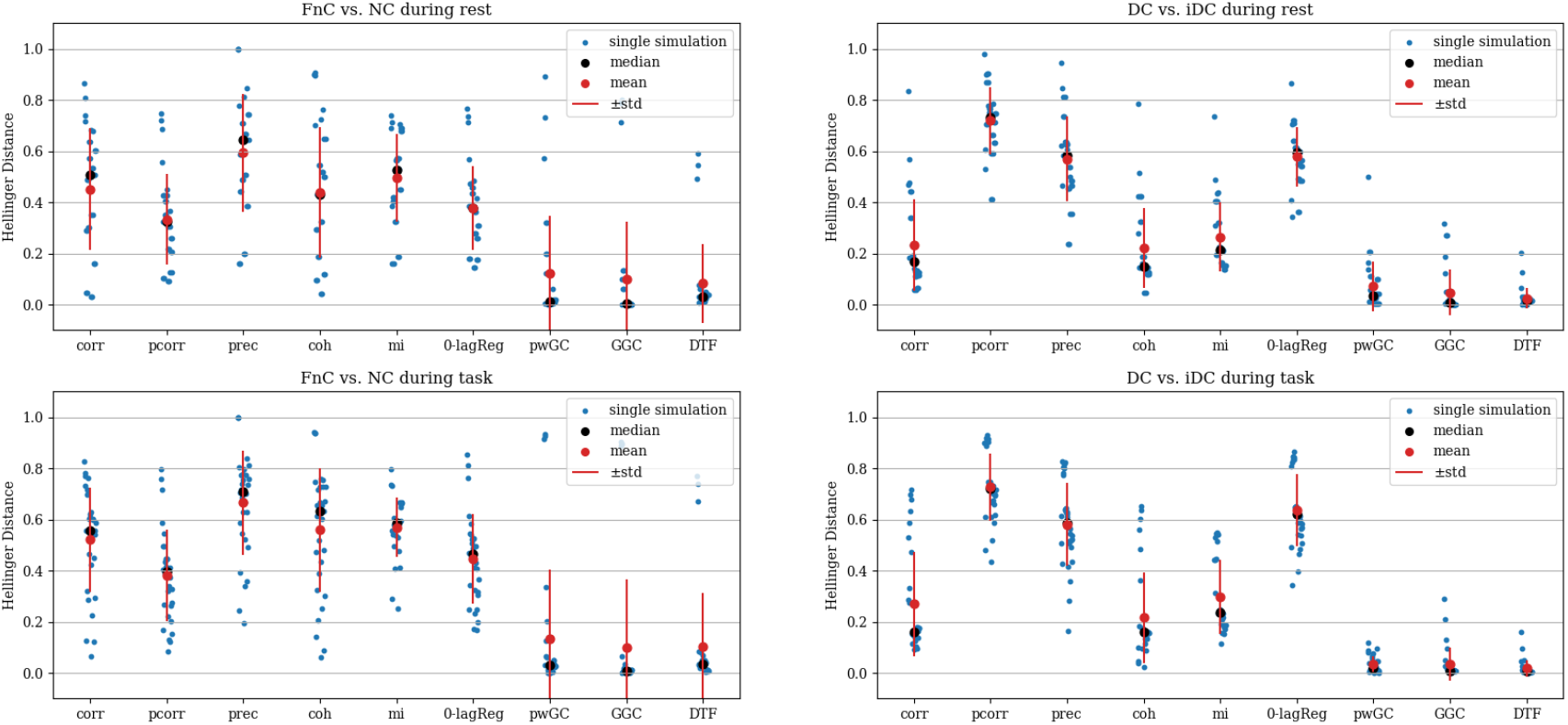
Hellinger distance between FnC vs. NC, and DC vs. iDC; summary over all simulations and methods during rest and task periods.

The implications of these findings are significant for advancing future research in functional connectivity. Researchers should carefully select FC methods based on the specific characteristics of the signals being analyzed, while also considering the impact of physiological artifacts. In many cases, multivariate methods offer an advantage by mitigating the effects of these artifacts, leading to more reliable connectivity estimates. Furthermore, optimizing signal quality and accounting for task-related influences are essential steps in ensuring accurate and robust connectivity measurements.

Future work should prioritize the development of artifact correction techniques for both neural signals and their corresponding functional connectivity measures. Additionally, exploring the effects of various experimental paradigms on connectivity measures will be critical for enhancing the accuracy and reliability of functional connectivity research. By refining these methods, researchers can better account for physiological artifacts and other confounding factors, ultimately improving the precision of connectivity assessments in both rest and task-based studies.

## References

[1] Zeki, S. and Shipp, S., 1988. The functional logic of cortical connections. Nature, 335(6188), pp.311–317.

[2] Friston, K.J., Frith, C.D., Liddle, P.F. and Frackowiak, R.S., 1993. Functional connectivity: the principal-component analysis of large (PET) data sets. Journal of Cerebral Blood Flow & Metabolism, 13(1), pp. 5–14.

[3] Friston, K.J., Frith, C.D. and Frackowiak, R.S.J., 1993. Time-dependent changes in effective connectivity measured with PET. Human Brain Mapping, 1(1), pp. 69–79.

[4] Etkin, A., Maron-Katz, A., Wu, W., Fonzo, G.A., Huemer, J., Vértes, P.E., Patenaude, B., Richiardi, J., Goodkind, M.S., Keller, C.J. and Ramos-Cejudo, J., 2019. Using fMRI connectivity to define a treatment-resistant form of post-traumatic stress disorder. Science translational medicine, 11(486), p. eaal3236.

[5] Lurie, D.J., Kessler, D., Bassett, D.S., Betzel, R.F., Breakspear, M., Kheilholz, S., Kucyi, A., Liégeois, R., Lindquist, M.A., McIntosh, A.R. and Poldrack, R.A., 2020. Questions and controversies in the study of time-varying functional connectivity in resting fMRI. Network neuroscience, 4(1), pp. 30–69.

[6] Foster, B.L., Rangarajan, V., Shirer, W.R. and Parvizi, J., 2015. Intrinsic and task-dependent coupling of neuronal population activity in human parietal cortex. Neuron, 86(2), pp. 578–590.

[7] Cole, M.W., Bassett, D.S., Power, J.D., Braver, T.S. and Petersen, S.E., 2014. Intrinsic and task-evoked network architectures of the human brain. Neuron, 83(1), pp. 238–251.

[8] Shine, J.M., Bissett, P.G., Bell, P.T., Koyejo, O., Balsters, J.H., Gorgolewski, K.J., Moodie, C.A. and Poldrack, R.A., 2016. The dynamics of functional brain networks: integrated network states during cognitive task performance. Neuron, 92(2), pp. 544–554.

[9] Ciuciu, P., Abry, P. and He, B.J., 2014. Interplay between functional connectivity and scale-free dynamics in intrinsic fMRI networks. Neuroimage, 95, pp.248-263.

[10] Van Mierlo, P., Papadopoulou, M., Carrette, E., Boon, P., Vandenberghe, S., Vonck, K. and Marinazzo, D., 2014. Functional brain connectivity from EEG in epilepsy: Seizure prediction and epileptogenic focus localization. Progress in neurobiology, 121, pp.19-35.

[11] Mitsis, G.D., Anastasiadou, M.N., Christodoulakis, M., Papathanasiou, E.S., Papacostas, S.S. and Hadjipapas, A., 2020. Functional brain networks of patients with epilepsy exhibit pronounced multiscale periodicities, which correlate with seizure onset. Human brain mapping, 41(8), pp. 2059–2076.

[12] Winterhalder, M., Schelter, B., Hesse, W., Schwab, K., Leistritz, L., Klan, D., Bauer, R., Timmer, J. and Witte, H., 2005. Comparison of linear signal processing techniques to infer directed interactions in multivariate neural systems. Signal processing, 85(11), pp. 2137–2160.

[13] Smith, S.M., Miller, K.L., Salimi-Khorshidi, G., Webster, M., Beckmann, C.F., Nichols, T.E., Ramsey, J.D. and Woolrich, M.W., 2011. Network modelling methods for FMRI. Neuroimage, 54(2), pp. 875–891.

[14] Askarinejad, S.E., Poline, J.B., and Mitsis, G.D., 2024, July. Investigation of the Effect of Physiological Artifacts on Task-based Functional Connectivity: A Simulation Study. In 2024 46th Annual International Conference of the IEEE Engineering in Medicine & Biology Society (EMBC) (pp. 1–5). IEEE.

[15] Sanz Leon, P., Knock, S.A., Woodman, M.M., Domide, L., Mersmann, J., McIntosh, A.R. and Jirsa, V., 2013. The Virtual Brain: a simulator of primate brain network dynamics. Frontiers in neuroinformatics, 7, p. 10.

[16] FitzHugh, R., 1961. Impulses and physiological states in theoretical models of nerve membrane. Biophysical journal, 1(6), pp. 445–466.

[17] Nagumo, J., Arimoto, S. and Yoshizawa, S., 1962. An active pulse transmission line simulating nerve axon. Proceedings of the IRE, 50(10), pp. 2061–2070.

[18] Stefanescu, R.A. and Jirsa, V.K., 2008. A low dimensional description of globally coupled heterogeneous neural networks of excitatory and inhibitory neurons. PLoS computational biology, 4(11), p. e1000219.

[19] Friston, K.J., Mechelli, A., Turner, R. and Price, C.J., 2000. Nonlinear responses in fMRI: the Balloon model, Volterra kernels, and other hemodynamics. NeuroImage, 12(4), pp. 466–477.

[20] Mann-Krzisnik, D. and Mitsis, G.D., 2022. Extracting electrophysiological correlates of functional magnetic resonance imaging data using the canonical polyadic decomposition. Human Brain Mapping, 43(13), pp. 4045–4073.

[21] Rundo, F., Conoci, S., Ortis, A. and Battiato, S., 2018. An advanced bio-inspired photoplethysmography (PPG) and ECG pattern recognition system for medical assessment. Sensors, 18(2), p. 405.

[22] Rassler, B., Schwerdtfeger, A., Aigner, C.S. and Pfurtscheller, G., 2018. “Switch-off” of respiratory sinus arrhythmia can occur in a minority of subjects during functional magnetic resonance imaging (fMRI). Frontiers in physiology, 9, p. 1688.

[23] Burrage, K., Burrage, P.M. and Tian, T., 2004. Numerical methods for strong solutions of stochastic differential equations: an overview. Proceedings of the Royal Society of London. Series A: Mathematical, Physical and Engineering Sciences, 460(2041), pp.373–402.

[24] Birn, R.M., Smith, M.A., Jones, T.B. and Bandettini, P.A., 2008. The respiration response function: the temporal dynamics of fMRI signal fluctuations related to changes in respiration. Neuroimage, 40(2), pp. 644–654.

[25] Chang, C., Cunningham, J.P. and Glover, G.H., 2009. Influence of heart rate on the BOLD signal: the cardiac response function. Neuroimage, 44(3), pp. 857–869.

[26] Kassinopoulos, M. and Mitsis, G.D., 2019. Identification of physiological response functions to correct for fluctuations in resting-state fMRI related to heart rate and respiration. Neuroimage, 202, p. 116150.

[27] Mclntosh, A.R. and Gonzalez-Lima, F., 1994. Structural equation modeling and its application to network analysis in functional brain imaging. Human brain mapping, 2(1-2), pp. 2–22.

[28] Marrelec, G., Krainik, A., Duffau, H., Pélégrini-Issac, M., Lehéricy, S., Doyon, J. and Benali, H., 2006. Partial correlation for functional brain interactivity investigation in functional MRI. Neuroimage, 32(1), pp. 228–237.

[29] Banerjee, O., Ghaoui, L.E., d’Aspremont, A. and Natsoulis, G., 2006, June. Convex optimization techniques for fitting sparse Gaussian graphical models. In Proceedings of the 23rd international conference on Machine learning (pp. 89–96).

[30] Friedman, J., Hastie, T. and Tibshirani, R., 2008. Sparse inverse covariance estimation with the graphical lasso. Biostatistics, 9(3), pp. 432–441.

[31] Welch, P., 1967. The use of fast Fourier transform for the estimation of power spectra: a method based on time averaging over short, modified periodograms. IEEE Transactions on audio and electroacoustics, 15(2), pp. 70–73.

[32] Kraskov, A., Stögbauer, H. and Grassberger, P., 2004. Estimating mutual information. Physical Review E—Statistical, Nonlinear, and Soft Matter Physics, 69(6), p. 066138.

[33] Shannon, C.E., 1948. A mathematical theory of communication. The Bell system technical journal, 27(3), pp. 379–423.

[34] Cover, T.M., 1999. Elements of information theory. John Wiley & Sons.

[35] Nieto-Castanon, A., 2020. Handbook of functional connectivity magnetic resonance imaging methods in CONN. Hilbert Press.

[36] Savanth, A.S., Pa, V., Nair, A.K. and Kutty, B.M., 2023. Differences in brain connectivity of meditators during assessing neurocognition via gamified experimental logic task: A machine learning approach. The Neuroradiology Journal, 36(3), pp. 305–314.

[37] Granger, C.W., 1969. Investigating causal relations by econometric models and cross-spectral methods. Econometrica: journal of the Econometric Society, pp.424–438.

[38] Geweke, J., 1982. Measurement of linear dependence and feedback between multiple time series. Journal of the American statistical association, 77(378), pp. 304–313.

[39] Akaike, H., 1974. A new look at the statistical model identification. IEEE transactions on automatic control, 19(6), pp. 716–723.

[40] Schwarz, G., 1978. Estimating the dimension of a model. The annals of statistics, pp. 461–464.

[41] Geweke, J.F., 1984. Measures of conditional linear dependence and feedback between time series. Journal of the American Statistical Association, 79(388), pp. 907–915.

[42] Kaminski, M.J. and Blinowska, K.J., 1991. A new method of the description of the information flow in the brain structures. Biological cybernetics, 65(3), pp. 203–210.

