## Supplementary Figures for "Investigation of the effect of physiological factors on resting-state and task-based functional connectivity"

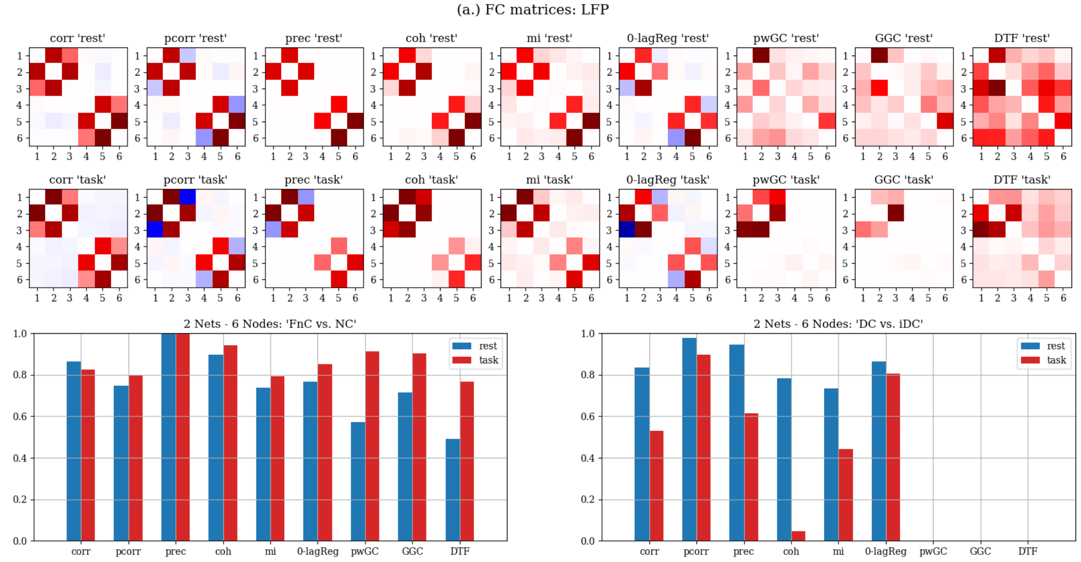


Figure S.1: Network of 6 nodes, LFP signals: (a) Rest- & task-FC matrices, (b) Hellinger distance: FnC vs NC, and DC vs iDC.

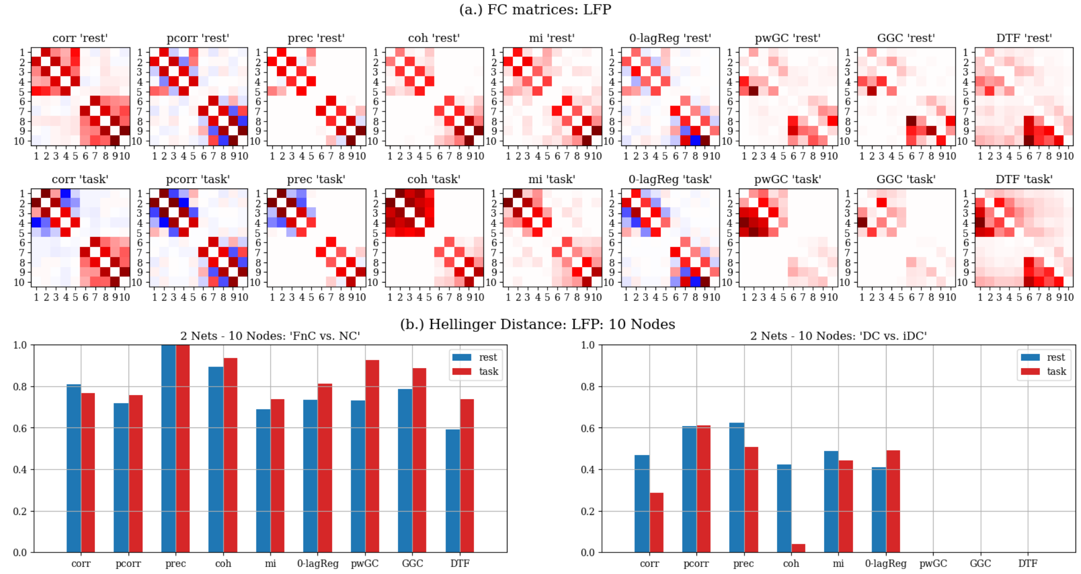


Figure S.2: Network of 10 nodes, LFP signals: (a) Rest- & task-FC matrices, (b) Hellinger distance: FnC vs NC, and DC vs iDC.

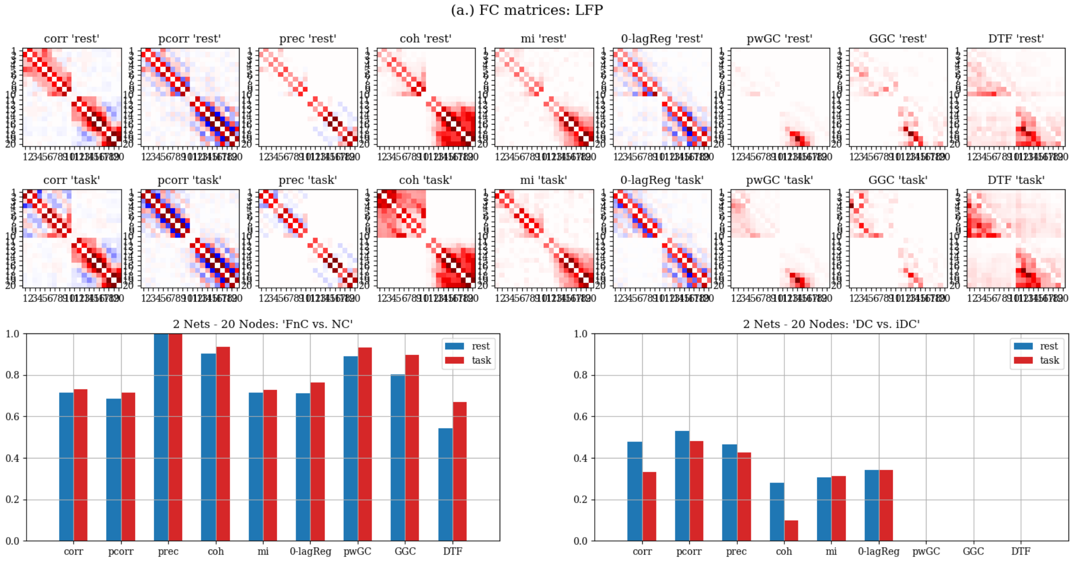


Figure S.3: Network of 20 nodes, LFP signals: (a) Rest- & task-FC matrices, (b) Hellinger distance: FnC vs NC, and DC vs iDC.

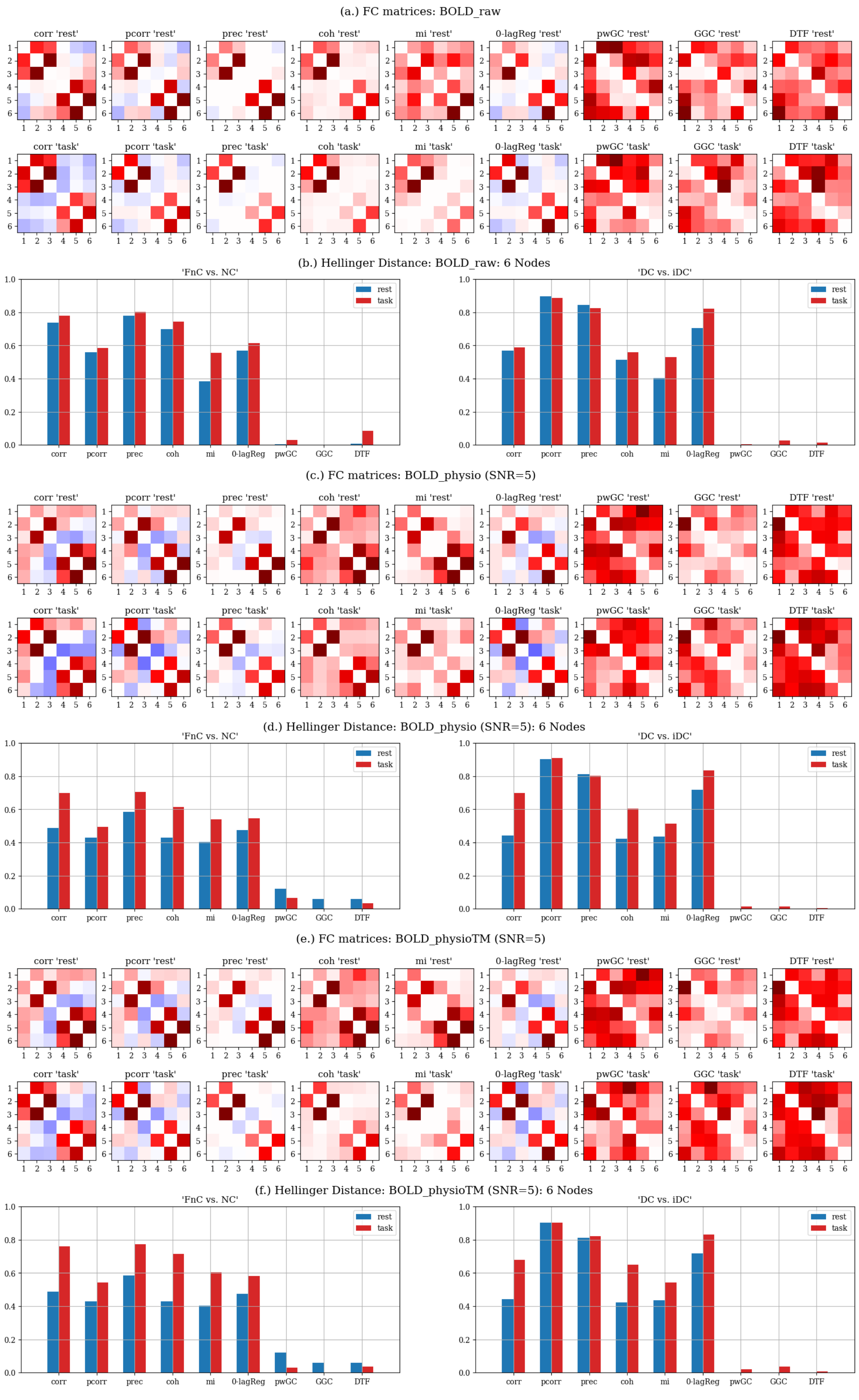


Figure S.4: Network of 6 nodes, BOLD signals: (a, c, e) Rest- & task-FC matrices, (b, d, f) Hellinger distance: FnC vs NC, and DC vs iDC (for BOLD_raw, BOLD_physio, and BOLD_physioTM datasets.)

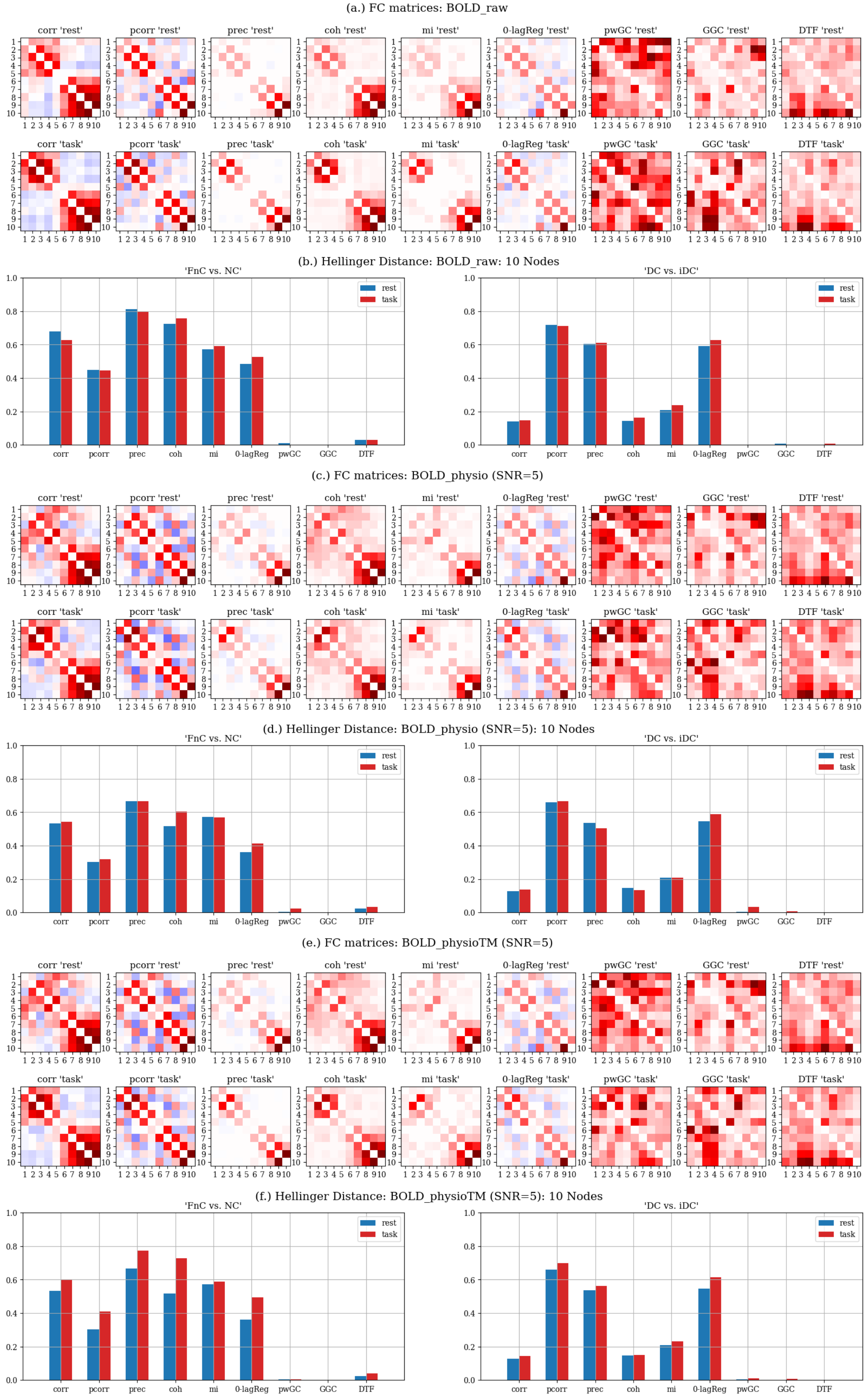


Figure S.5: Network of 10 nodes, BOLD signals: (a, c, e) Rest- & task-FC matrices, (b, d, f) Hellinger distance: FnC vs NC, and DC vs iDC (for BOLD_raw, BOLD_physio, and BOLD_physioTM datasets.)

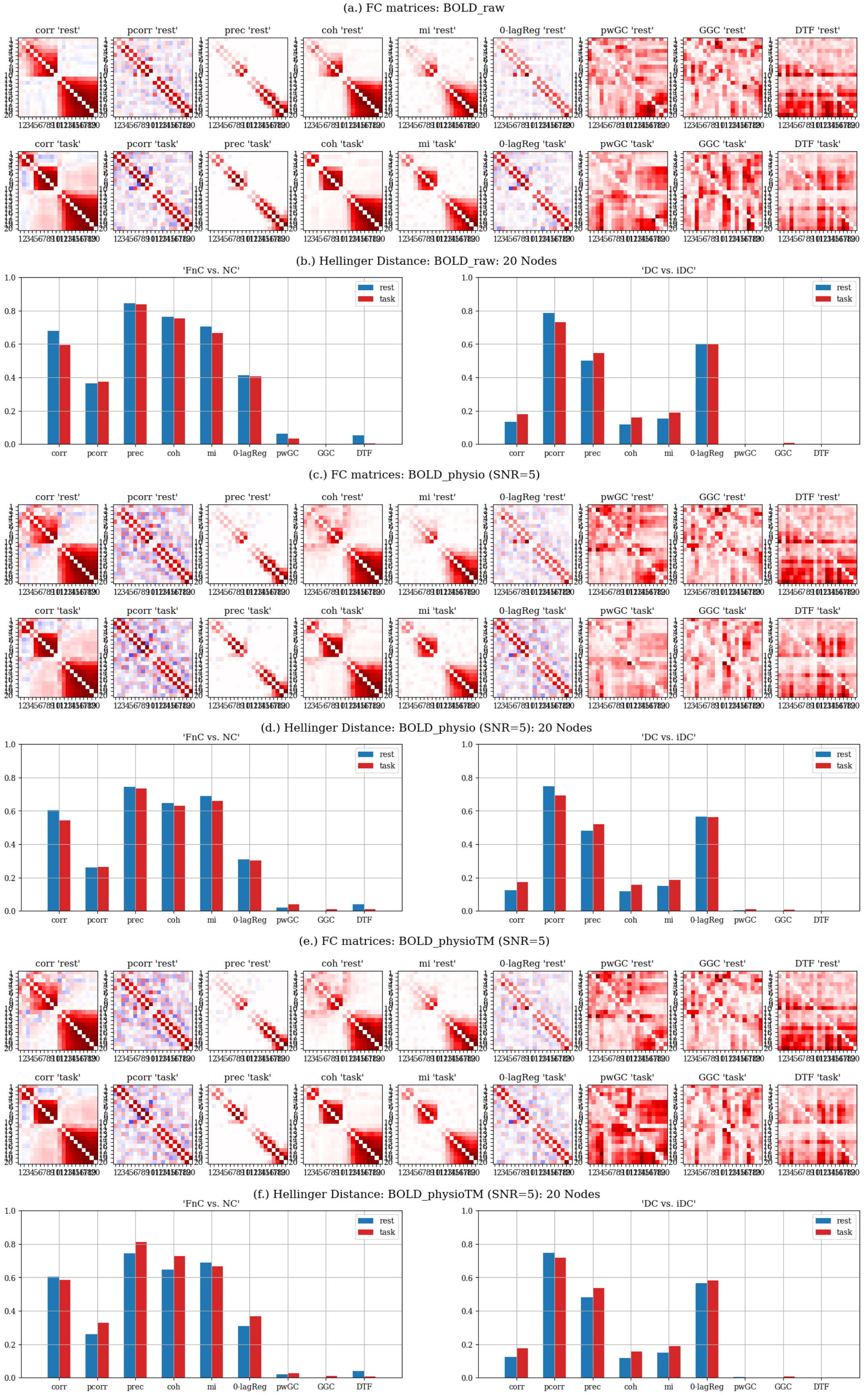


Figure S.6: Network of 20 nodes, BOLD signals: (a, c, e) Rest- & task-FC matrices, (b, d, f) Hellinger distance: FnC vs NC, and DC vs iDC (for BOLD_raw, BOLD_physio, and BOLD_physioTM datasets.)

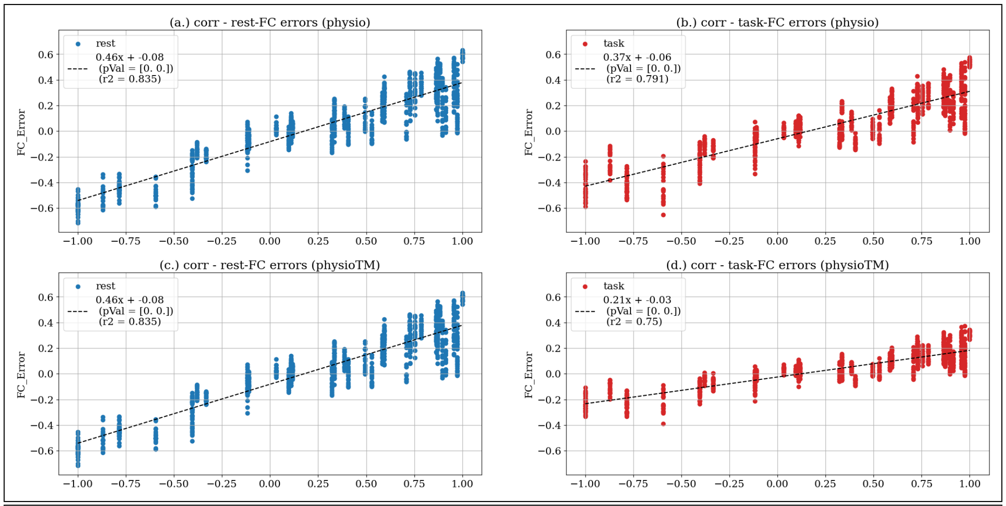
(a.) corr - rest-FC errors (physio)
rest
0.46x + -0.08
(pVal = [0. 0.])
(r2 = 0.835)
FC_Error
(b.) corr - task-FC errors (physio)
task
0.37x + -0.06
(pVal = [0. 0.])
(r2 = 0.791)
FC_Error
(c.) corr - rest-FC errors (physioTM)
rest
0.46x + -0.08
(pVal = [0. 0.])
(r2 = 0.835)
FC_Error
(d.) corr - task-FC errors (physioTM)
task
0.21x + -0.03
(pVal = [0. 0.])
(r2 = 0.75)
FC_Error

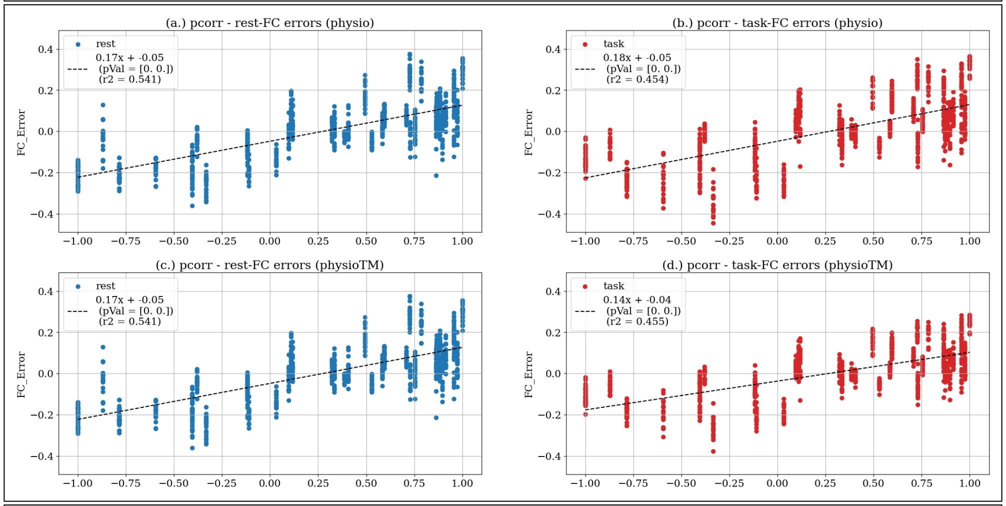
(a.) pcorr - rest-FC errors (physio)
rest
0.17x + -0.05
(pVal = [0. 0.])
(r2 = 0.541)
FC_Error
(b.) pcorr - task-FC errors (physio)
task
0.18x + -0.05
(pVal = [0. 0.])
(r2 = 0.454)
FC_Error
(c.) pcorr - rest-FC errors (physioTM)
rest
0.17x + -0.05
(pVal = [0. 0.])
(r2 = 0.541)
FC_Error
(d.) pcorr - task-FC errors (physioTM)
task
0.14x + -0.04
(pVal = [0. 0.])
(r2 = 0.455)
FC_Error

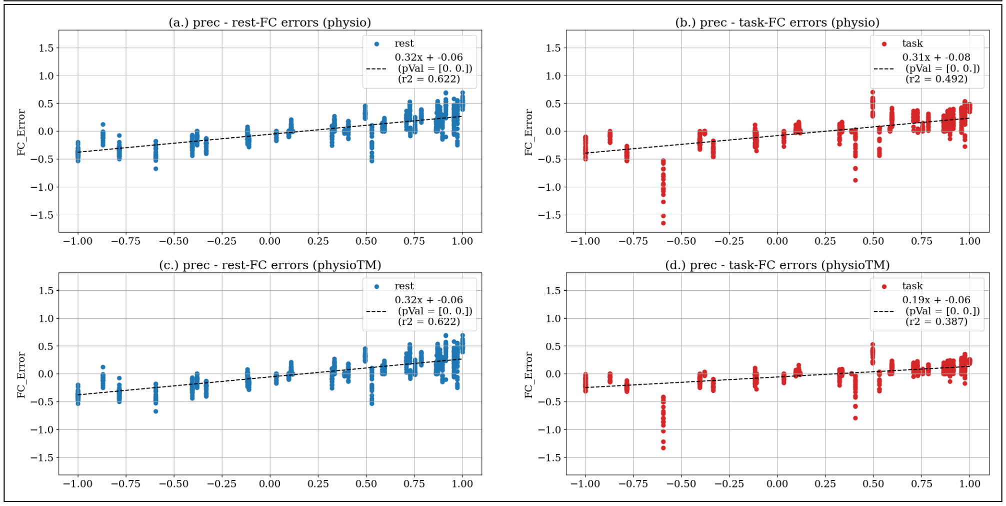
(a.) prec - rest-FC errors (physio)
rest
0.32x + -0.06
(pVal = [0. 0.])
(r2 = 0.622)
FC_Error
(b.) prec - task-FC errors (physio)
task
0.31x + -0.08
(pVal = [0. 0.])
(r2 = 0.492)
FC_Error
(c.) prec - rest-FC errors (physioTM)
rest
0.32x + -0.06
(pVal = [0. 0.])
(r2 = 0.622)
FC_Error
(d.) prec - task-FC errors (physioTM)
task
0.19x + -0.06
(pVal = [0. 0.])
(r2 = 0.387)
FC_Error

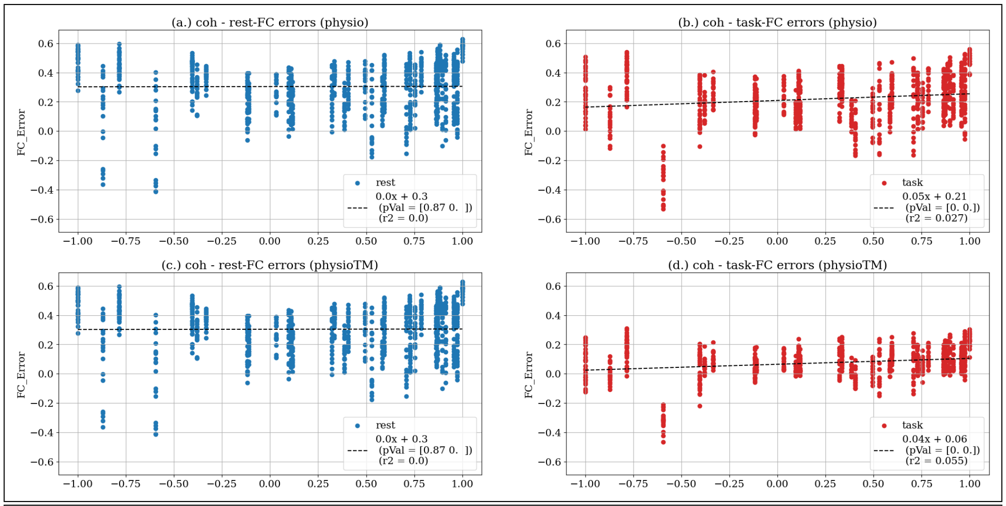
(a.) coh - rest-FC errors (physio)
FC_Error
rest
0.0x + 0.3
(pVal = [0.87 0. ])
(r2 = 0.0)
(b.) coh - task-FC errors (physio)
FC_Error
task
0.05x + 0.21
(pVal = [0. 0.])
(r2 = 0.027)
(c.) coh - rest-FC errors (physioTM)
FC_Error
rest
0.0x + 0.3
(pVal = [0.87 0. ])
(r2 = 0.0)
(d.) coh - task-FC errors (physioTM)
FC_Error
task
0.04x + 0.06
(pVal = [0. 0.])
(r2 = 0.055)

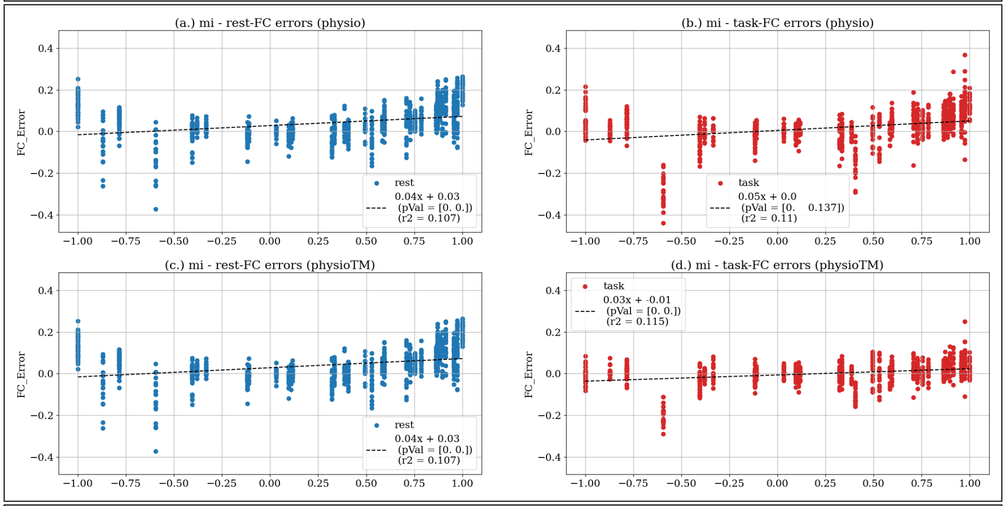
(a.) mi - rest-FC errors (physio)
FC_Error
rest
0.04x + 0.03
(pVal = [0. 0.])
(r2 = 0.107)
(b.) mi - task-FC errors (physio)
FC_Error
task
0.05x + 0.0
(pVal = [0. 0.137])
(r2 = 0.11)
(c.) mi - rest-FC errors (physioTM)
FC_Error
rest
0.04x + 0.03
(pVal = [0. 0.])
(r2 = 0.107)
(d.) mi - task-FC errors (physioTM)
FC_Error
task
0.03x + -0.01
(pVal = [0. 0.])
(r2 = 0.115)

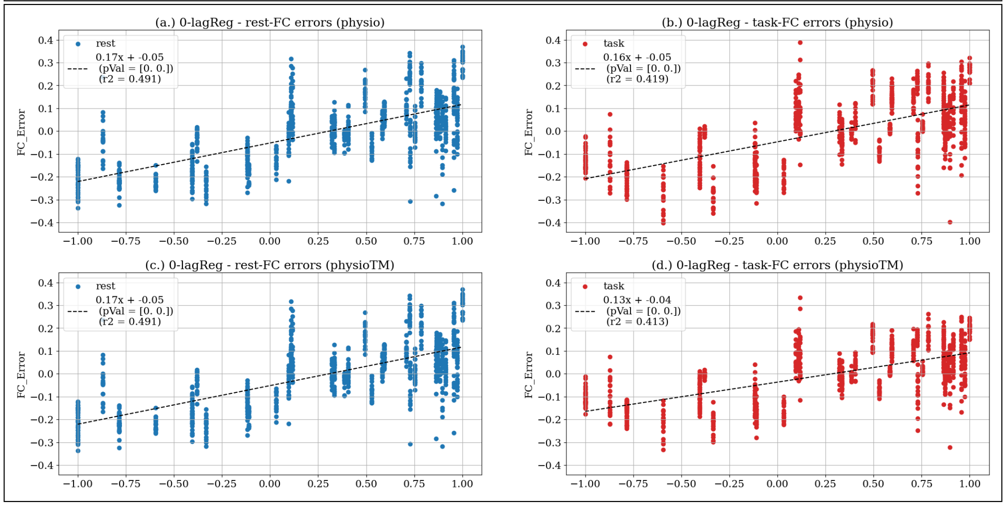
(a.) 0-lagReg - rest-FC errors (physio)
FC_Error
rest
0.17x + -0.05
(pVal = [0. 0.])
(r2 = 0.491)
(b.) 0-lagReg - task-FC errors (physio)
FC_Error
task
0.16x + -0.05
(pVal = [0. 0.])
(r2 = 0.419)
(c.) 0-lagReg - rest-FC errors (physioTM)
FC_Error
rest
0.17x + -0.05
(pVal = [0. 0.])
(r2 = 0.491)
(d.) 0-lagReg - task-FC errors (physioTM)
FC_Error
task
0.13x + -0.04
(pVal = [0. 0.])
(r2 = 0.413)

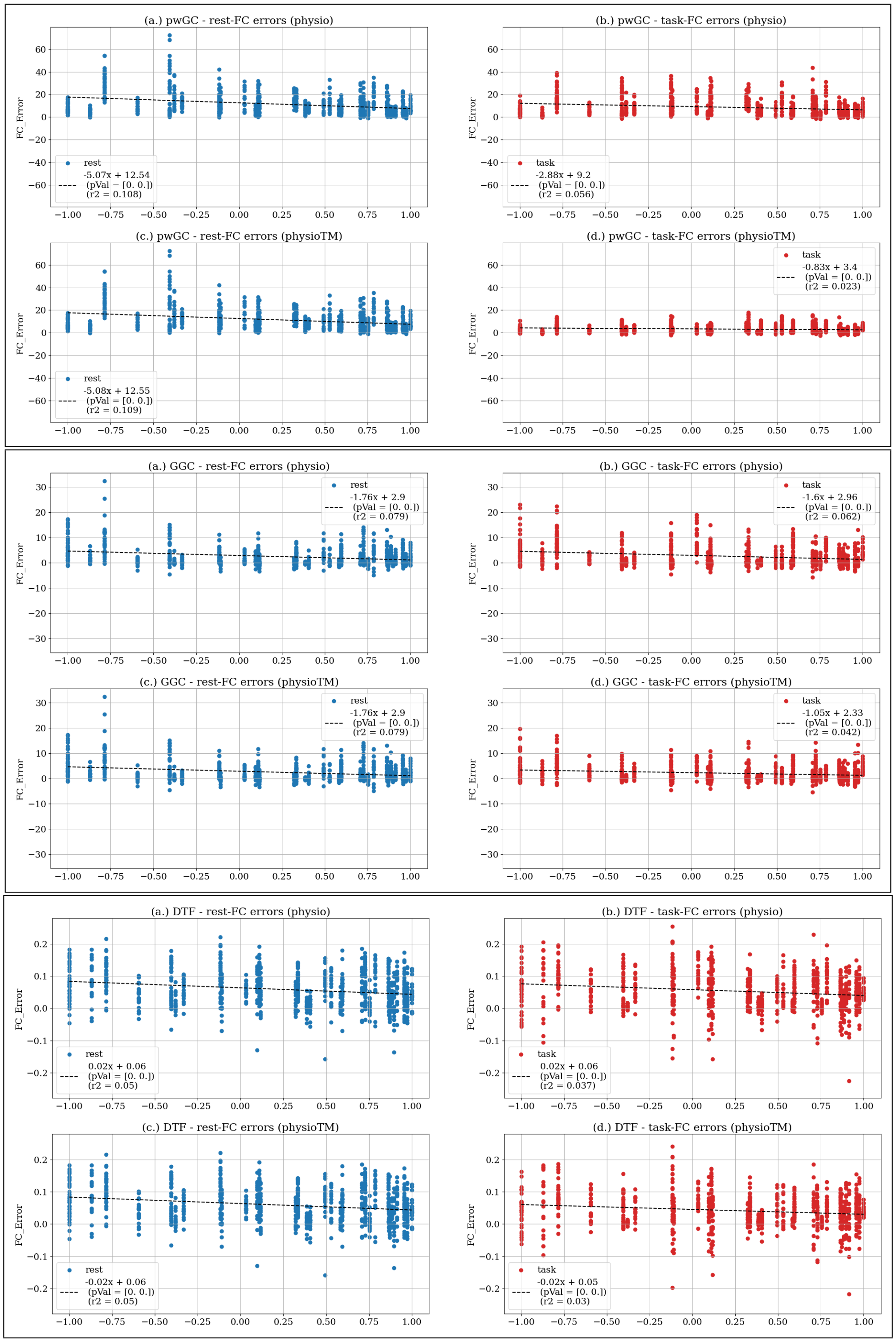


Figure S.9: Errors induced by physiological artifacts to rest- & task-FC values vs. correlation of CRFs: over all BOLD simulations

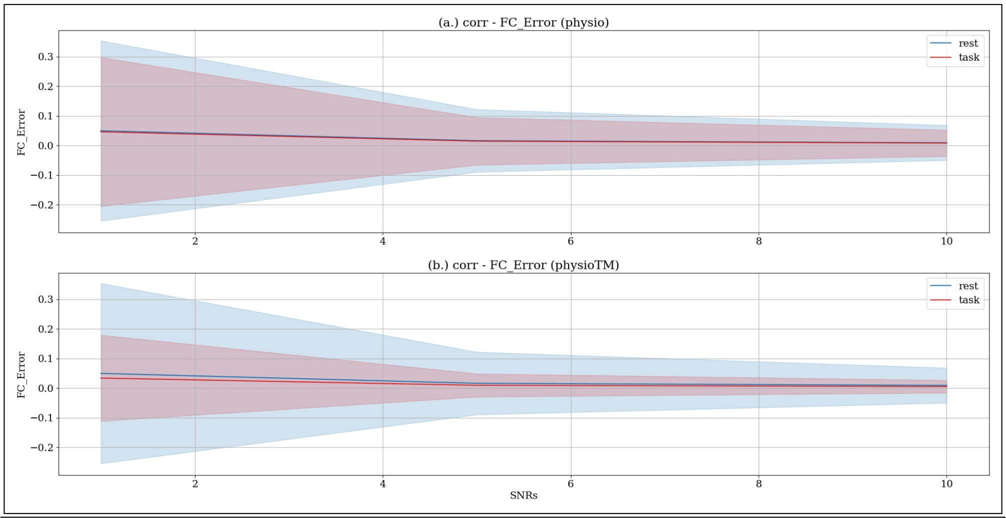
(a.) corr - FC_Error (physio)
FC_Error
rest
task
(b.) corr - FC_Error (physioTM)
FC_Error
rest
task
SNRs

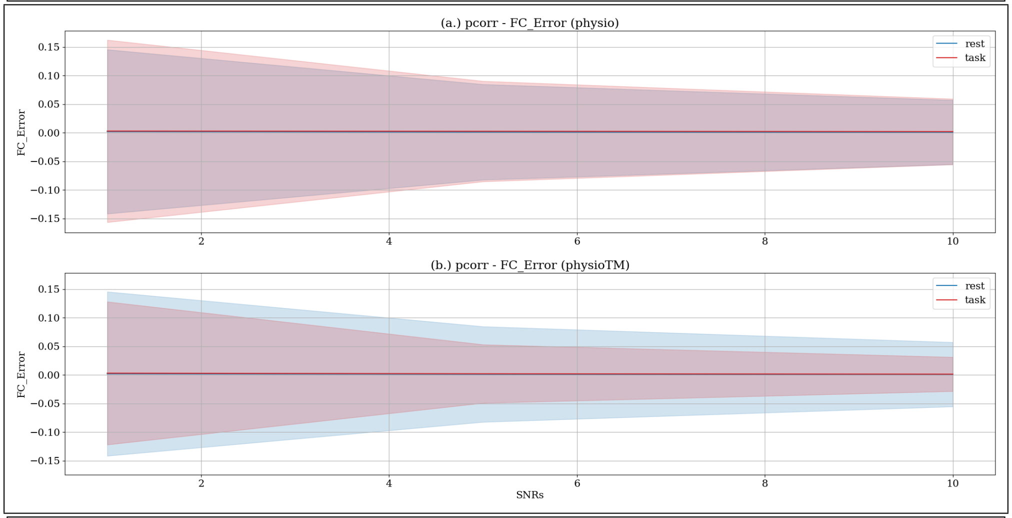
(a.) pcorr - FC_Error (physio)
FC_Error
rest
task
(b.) pcorr - FC_Error (physioTM)
FC_Error
rest
task
SNRs

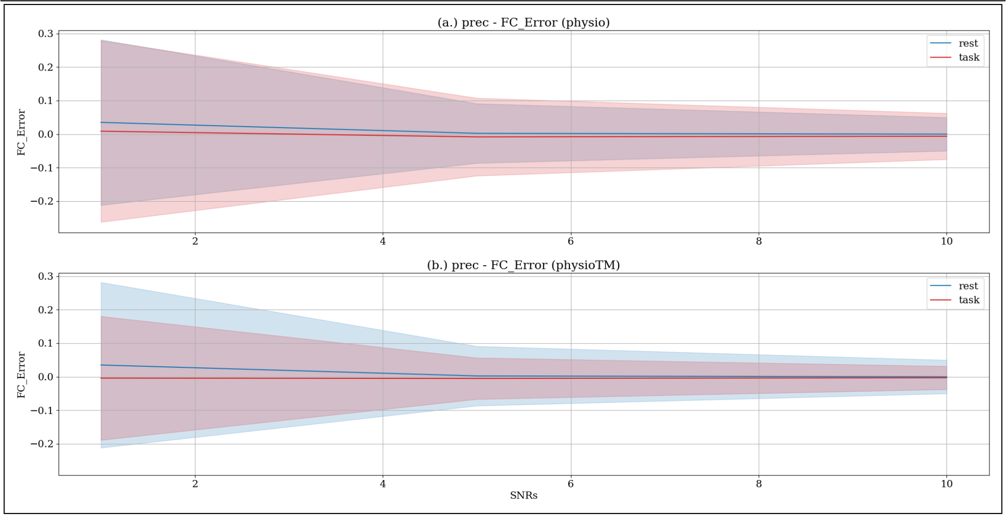
(a.) prec - FC_Error (physio)
FC_Error
rest
task
(b.) prec - FC_Error (physioTM)
FC_Error
rest
task
SNRs

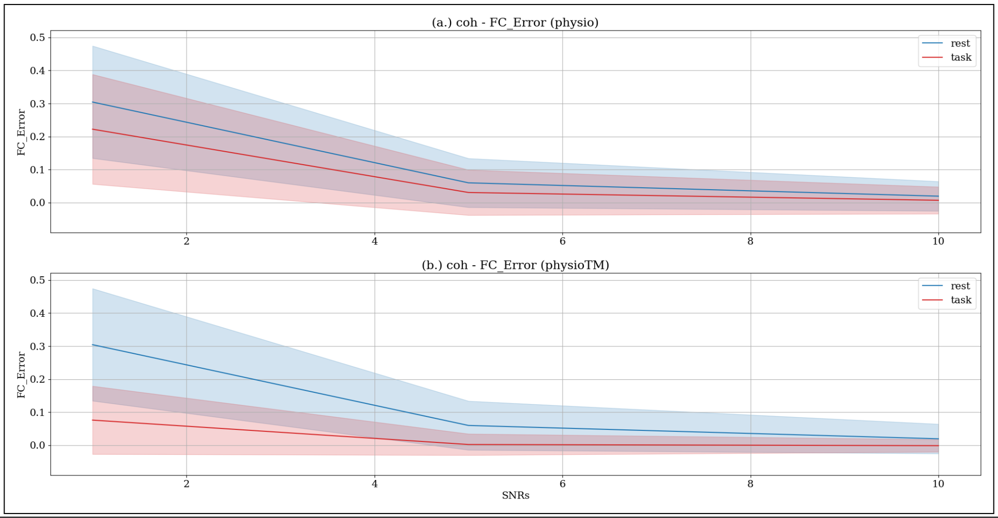
(a.) coh - FC_Error (physio)
(b.) coh - FC_Error (physioTM)
rest
task
FC_Error
SNRs

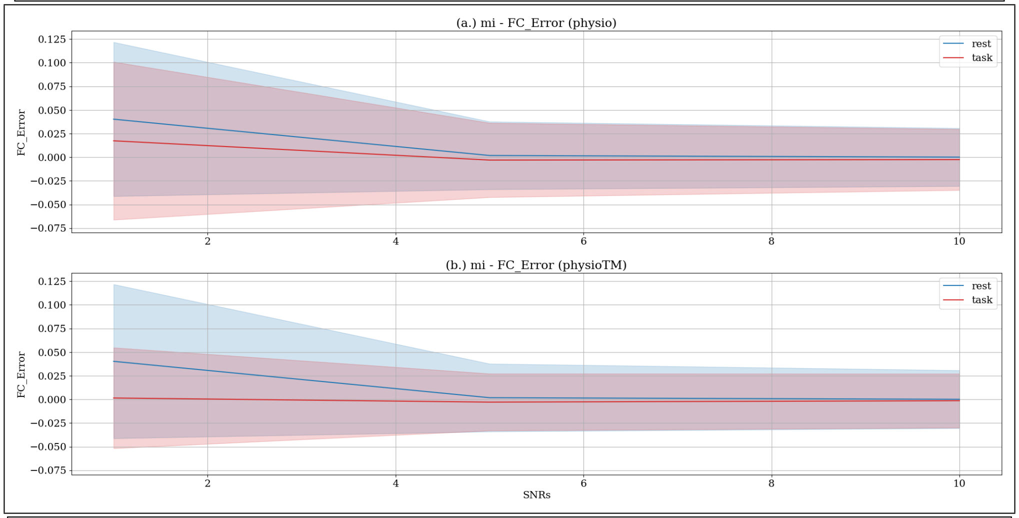
(a.) mi - FC_Error (physio)
(b.) mi - FC_Error (physioTM)
rest
task
FC_Error
SNRs

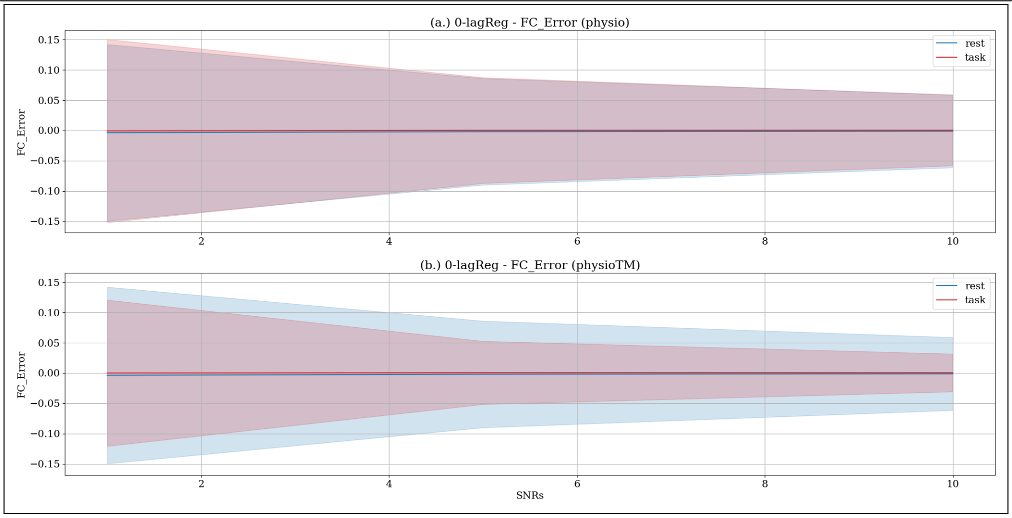
(a.) 0-lagReg - FC_Error (physio)
(b.) 0-lagReg - FC_Error (physioTM)
rest
task
FC_Error
SNRs

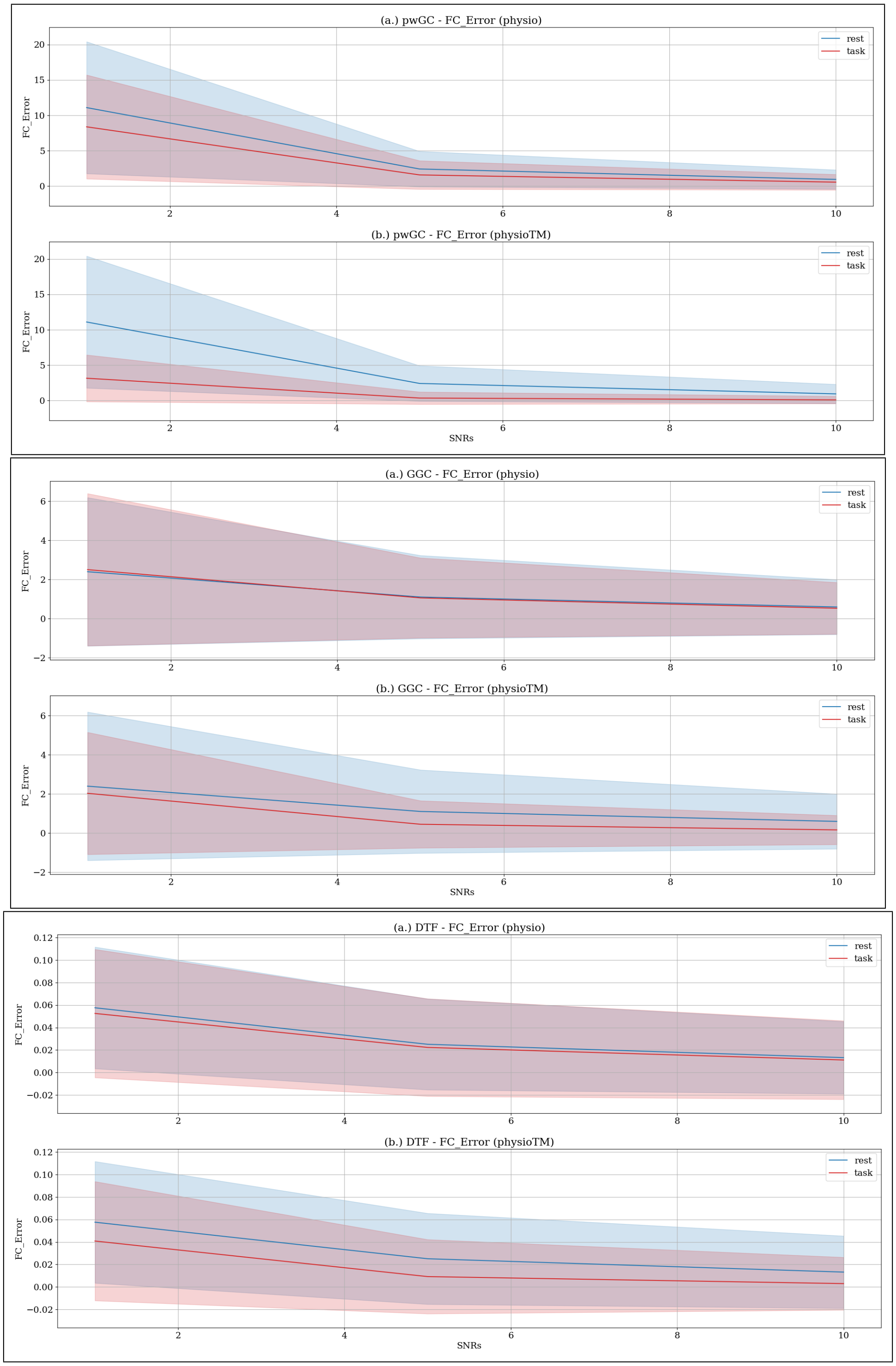


Figure S.12: Mean and standard deviation band of the errors induced by physiological artifacts to rest- & task-FC values across various SNRs: over all BOLD simulations

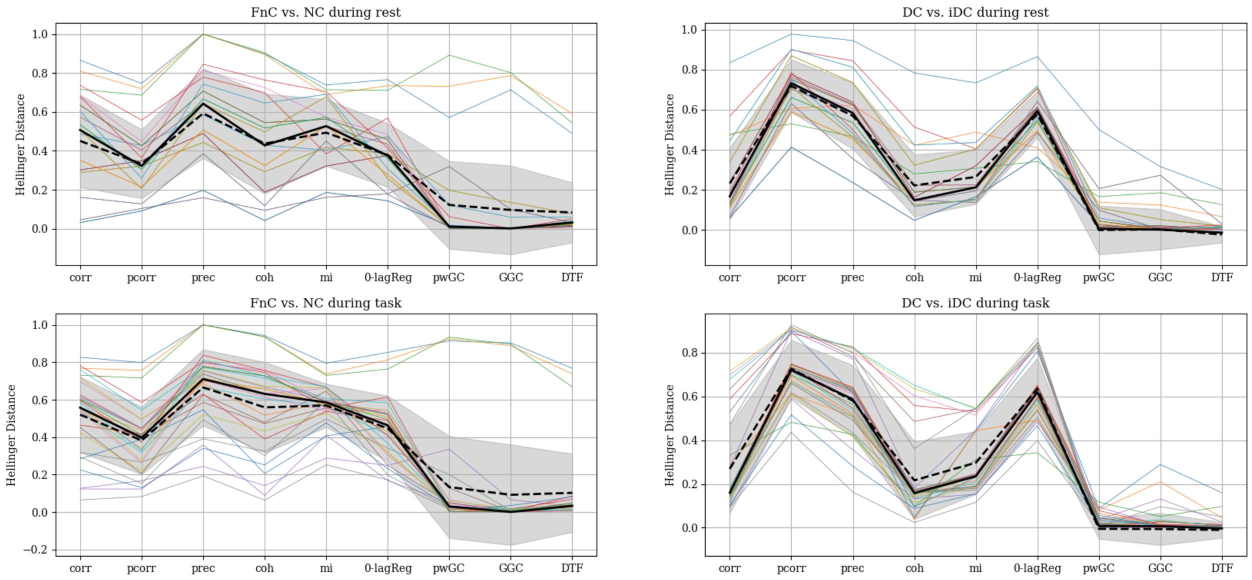


Figure S.13: Hellinger distance between FnC vs. NC, and DC vs. iDC; summary over all simulations and methods during rest and task periods. Each line represents a single simulation, the thick black line illustrates the median of the results, the dashed line shows the mean and the gray area is the ±std interval.
